# Pan-pediatric brain tumor microRNA networks define immune microenvironment states and responses to CAR T-cell therapy

**DOI:** 10.64898/2026.10.21.760687

**Authors:** Kaleem Coleman, Ryan Corbett, Zhongzhen Yi, Bicna Song, Alex Sickler, Aylar Babaei, Jo Lynne Rokita, Dalia Haydar

## Abstract

Pediatric central nervous system (CNS) tumors remain a leading cause of cancer-related mortality in children, yet molecular programs that shape their immune microenvironments and response to immunotherapy remain poorly defined. Because microRNAs (miRNAs) coordinately regulate gene expression across multiple pathways, we hypothesized that miRNA programs may capture broader immune states than individual immune-regulatory molecules. We integrated small RNA and bulk transcriptomic profiles from diffuse midline glioma, medulloblastoma, ependymoma, and low-grade glioma to identify miRNA programs associated with tumor immune states. Across tumor histologies, dysregulated miRNAs were associated with T-cell, natural killer cell, antigen-presentation, and myeloid programs. Diffuse midline glioma exhibited a distinct miRNA program associated with reduced lymphocyte signatures and increased suppressive myeloid features. Longitudinal profiling of immunocompetent glioma models during B7-H3 CAR T-cell therapy revealed dynamic remodeling of miRNA and immune transcriptional programs. Human tumor-associated miRNAs were significantly enriched among miRNAs associated with CAR T response *in vivo*. Functional interrogation of miR-149 demonstrated enhanced CAR T-cell expansion and sustained antitumor activity during repeated antigen stimulation. Together, these data define miRNA programs associated with immune states across pediatric CNS tumors and identify miR-149 as a candidate regulator of CAR T-cell function, providing a genomic framework for developing miRNA-directed strategies to improve CAR T-cell therapy.

**Importance of Study:** Pediatric CNS tumors exhibit distinct immunomodulatory microRNA signatures that are associated with an immunosuppressive tumor microenvironment and are altered following B7-H3 CAR T-cell therapy. Identification of therapy-responsive microRNAs conserved across preclinical models and patient samples provides potential targets for simultaneously modulating multiple suppressive pathways and improving immunotherapy outcomes in pediatric brain tumors.

## Introduction

Pediatric central nervous system (CNS) tumors are a leading cause of cancer-related death in children and encompass biologically diverse malignancies, including diffuse midline glioma (DMG), high-grade glioma (HGG), and medulloblastoma (MB) (1). DMG, historically referred to as diffuse intrinsic pontine glioma (DIPG) when located in the pons, remains among the most lethal pediatric CNS tumors, with a 5-year survival rate below 2% (2). MB and HGG similarly comprise multiple molecular and clinical subtypes with substantial inter- and intratumoral heterogeneity (3). Although advances in genomic and epigenomic profiling have refined the classification of pediatric brain tumors, this improved molecular understanding has not consistently translated into durable therapeutic responses (3). New strategies are therefore needed to address not only tumor-intrinsic heterogeneity but also the broader biological networks that support treatment resistance.

Chimeric antigen receptor (CAR) T-cell therapy has transformed the treatment of selected hematologic malignancies but has only achieved limited and often non-durable activity in solid tumors, including pediatric brain tumors (4). CAR T cells targeting tumor-associated antigens such as B7-H3 demonstrate potent activity in preclinical models, supporting their therapeutic potential across pediatric CNS malignancies (5). However, early clinical studies have largely produced partial, heterogeneous, or transient responses (6). Multiple interconnected barriers contribute to these limitations, including antigen heterogeneity, inadequate trafficking and tumor penetration, T-cell exhaustion, and limited persistence. Importantly, many of these barriers are imposed or reinforced by the tumor microenvironment (TME), which can restrict CAR T-cell infiltration, activation, and durability even when the targeted antigen is present (7).

The pediatric brain tumor microenvironment is a dynamic ecosystem shaped by interactions among malignant cells, infiltrating and tissue-resident immune populations, stromal elements, and neural-lineage cells (8). Immune and neuroregulatory signals within this ecosystem can promote suppressive myeloid states, impair T-cell function, and enable tumors to adapt under therapeutic pressure (7,8). We propose that the pediatric brain TME is organized as an interconnected regulatory network capable of transitioning between distinct immune states rather than as a static collection of individual suppressive factors. These states may differ across tumor types and molecular subtypes while also sharing common programs that promote immune suppression and treatment resistance. Identifying the molecular networks that govern these dynamic states is therefore critical for understanding why some tumors respond to immunotherapy while others remain resistant. This framework also suggests that disrupting individual suppressive pathways may be insufficient when resistance is maintained by broader, interconnected regulatory programs.

Most strategies designed to improve CAR T-cell activity target a single antigen, receptor, checkpoint, cytokine, signaling pathway, or suppressive cell population (9,10). Although such approaches can disrupt individual resistance mechanisms, they may be insufficient to durably remodel a TME maintained by redundant and interconnected immune, tumor-intrinsic, and neuroregulatory pathways. Inhibition of one pathway can leave parallel mechanisms intact or induce compensatory programs that preserve immune suppression. Improving CAR T-cell efficacy may therefore require immune-adjuvant strategies that act across multiple pathways rather than further optimization of a single target. This creates a need to identify systems-level regulators that can both predict changes in TME state and coordinate broader remodeling of the tumor ecosystem. We hypothesized that effective remodeling of pediatric brain tumor microenvironments requires coordinated regulation of multiple suppressive pathways rather than modulation of individual immune mechanisms, and that microRNAs may function as key regulators of these broader immune programs.

MicroRNAs (miRNAs) are well positioned to fulfill this role. These conserved, non-coding RNAs act as post-transcriptional regulators of gene expression, with individual miRNAs capable of regulating numerous transcripts across interconnected signaling networks (11). Unlike strategies directed against a single molecular target, miRNAs can coordinate programs involved in immune-cell differentiation, myeloid polarization, T-cell activation and persistence, neuronal signaling, and tumor progression (12). Their expression patterns may therefore define broader tumor ecosystem states rather than reflect isolated molecular alterations (13,14). Moreover, because dysregulated miRNAs can regulate multiple components of an immunosuppressive network, they may serve not only as biomarkers of TME composition and therapeutic response but also as actionable targets for simultaneously modulating several mechanisms of immune suppression (14).

Despite this potential, the systems-level contribution of miRNAs to the pediatric brain tumor TME remains poorly defined. Most studies have focused on individual miRNAs, isolated pathways, or single tumor types, limiting the identification of shared regulatory programs across biologically distinct pediatric CNS tumors. It remains unclear whether pan-tumor miRNA programs exist across DMG, MB, HGG, and other pediatric brain tumor subtypes, or whether such programs are associated with common immune and neuronal TME states. The relationship between miRNA dysregulation and transcriptome-defined TME signatures is also insufficiently understood. Furthermore, few studies have examined how these miRNA programs change in response to CAR T-cell therapy or whether therapy-responsive miRNAs are conserved between patient tumors and immunocompetent preclinical models (15,16). This limited integration of patient biology, TME state, and therapeutic response has constrained the development of predictive miRNA signatures and miRNA-directed strategies for immunotherapy.

In this study, we integrated small RNA sequencing with bulk transcriptomic profiling across multiple pediatric brain tumor histologies and molecular subtypes, including DMG, MB, and HGG, to identify dysregulated miRNAs associated with immune and neuronal features of the TME. We correlated tumor-associated miRNA expression with transcriptome-derived TME signatures to define miRNA programs associated with distinct immunoregulatory states across pediatric brain tumors. We then interrogated miRNA remodeling following B7-H3 CAR T-cell therapy in immunocompetent glioma models and examined the conservation of therapy-associated miRNAs between preclinical tumors and patient-derived DMG samples. Finally, we functionally evaluated one candidate miRNA to determine its impact on CAR T-cell activity. By combining pan-tumor patient profiling, integrated miRNA and transcriptomic analyses, immunocompetent therapeutic models, cross-species validation, and functional interrogation, this study identifies miRNA programs that reflect TME state and response to CAR T-cell therapy. These findings provide a foundation for developing miRNA-based biomarkers of TME remodeling and multi-pathway strategies to overcome immune suppression and improve CAR T-cell therapy for pediatric brain tumors.

## Results

### Pediatric CNS tumors harbor histology-specific and shared miRNA programs

To define miRNA programs associated with pediatric CNS tumors, we performed small RNA sequencing on 21 post-mortem tumors representing diffuse midline glioma (DMG; n = 11), medulloblastoma (MB; n = 5), ependymoma (EPN; n = 3), and low-grade glioma (LGG; n = 2). The cohort also included tumor-adjacent brain tissue from 11 patients, comprising five DMG, four MB, and two EPN cases, and normal brain tissue from five pediatric donors without cancer (**Figure 1A; Table S1**).

**Figure 1.**
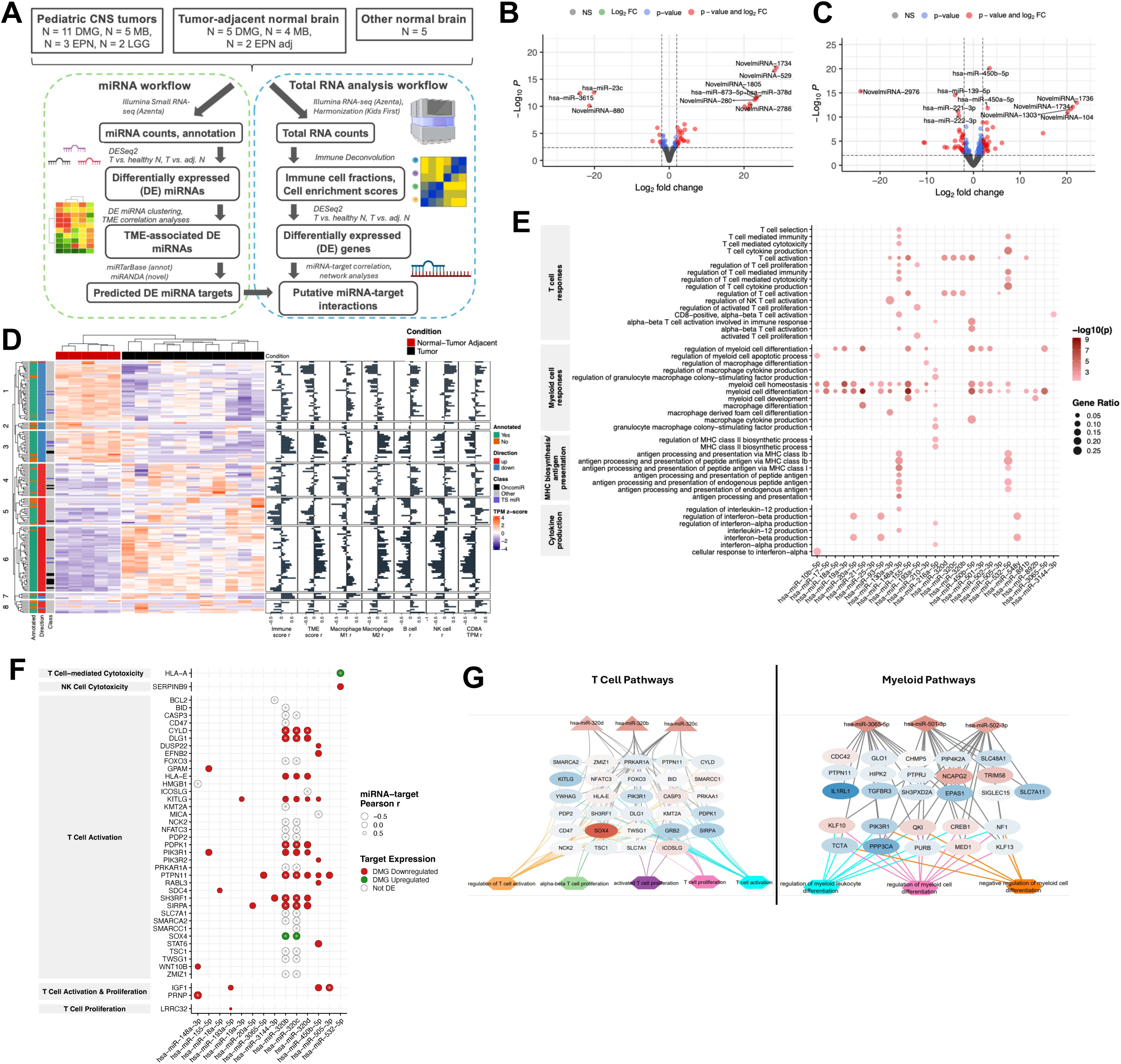
DMG-associated miRNA programs are linked to suppressive immune microenvironment states. **(A)** Study design and integrated small RNA-seq and bulk RNA-seq workflow for pediatric CNS tumor and non-tumor brain samples. **(B-C)** Volcano plots showing differential miRNA expression in DMG relative to healthy pediatric brain (**B**; n = 11 DMG, n = 5 healthy brain) and tumor-adjacent brain (**C**; n = 5/group). Dashed lines indicate FDR < 0.05 and |log fold change| > 2. **(D)** Hierarchical clustering of differentially expressed miRNAs in DMG and tumor-adjacent brain tissue. Adjacent bar plots show Pearson correlations between cluster expression and inferred immune-cell abundance, immune scores, tumor microenvironment (TME) scores, and *CD8A* expression. **(E)** Gene Ontology enrichment analysis of predicted targets of cluster 6 miRNAs showing enrichment of immune-related biological processes. **(F)** Network of significant associations between cluster 6 miRNAs and target genes involved in T-cell-related pathways. Interactions were supported by inverse miRNA-target correlations and/or opposing differential expression patterns. Asterisks denote Pearson correlation *p* < 0.05. **(G)** Simplified miRNA-target-pathway network derived from cluster 6. Triangles indicate miRNAs, circles indicate target genes, and hexagons indicate enriched biological pathways. Node colors represent differential expression (red, upregulated; blue, downregulated).

Principal-component analysis of both miRNA and mRNA expression profiles separated pediatric CNS tumors from tumor-adjacent and healthy brain tissues, revealing broad tumor-associated differences in the small RNA and transcriptional landscapes (**Figure S1**). We therefore examined miRNA dysregulation within individual histologies and integrated these profiles with matched transcriptomic features of the TME.

### DMGs harbor a miRNA program associated with a suppressive immune TME

We first asked whether DMG contains coordinated miRNA programs associated with tumor biology and immune regulation. Differential expression analysis comparing DMG with tumor-adjacent or healthy pediatric brain identified 97 significantly upregulated and 73 significantly downregulated miRNAs (|log fold change| > 0, Wald test false discovery rate [FDR] < 0.05; **Figures 1B, 1C, S2A, and S2B; Tables S2A and S2B**). Unsupervised clustering of these differentially expressed miRNAs separated DMG from non-tumor brain and identified eight expression clusters, indicating that tumor-associated miRNAs are organized into distinct regulatory programs rather than a uniform pattern of dysregulation (**Figure 1D; Table S3A**).

To determine whether these programs contained miRNAs with established roles in tumor biology, we compared the differentially expressed miRNAs with a curated set of experimentally validated oncogenic miRNAs and tumor-suppressive miRNAs (**Table S4**). Thirteen validated oncogenic miRNAs and ten tumor-suppressive miRNAs were differentially expressed in DMG. The direction of dysregulation was concordant with their reported functions for 11 of 13 oncogenic miRNAs and seven of ten tumor-suppressive miRNAs. Cluster 6 was particularly notable: it contained 46 miRNAs, including 45 upregulated miRNAs and one downregulated miRNA, and had the highest proportion of validated oncogenic miRNAs among the eight clusters (8 of 46, 17.4%). Thus, cluster 6 emerged as a prominent DMG-associated program with both oncogenic and potentially immunoregulatory features.

We next asked whether the DMG-associated miRNA clusters corresponded to distinct immune states within the TME. Cell-type deconvolution of matched bulk RNA-seq data revealed substantial heterogeneity in inferred immune composition across DMG tumors (**Figure S3**). Integration of the deconvolution results with miRNA expression demonstrated cluster-specific relationships with inferred immune populations and composite TME scores (**Figure 1D**).

Among the identified programs, cluster 6 showed the strongest association with suppressive immune features. Cluster 6 expression was inversely associated with inferred B-cell abundance (mean Pearson r = −0.32, range = −0.83-0.20), NK-cell abundance (mean Pearson r = −0.34, range = −0.94-0.40), and *CD8A* expression (mean Pearson r = −0.31, range = −0.76-0.22), but positively associated with inferred M2-like macrophage abundance (mean Pearson r = 0.32, range = −0.58-0.40). These findings identify cluster 6 as a DMG-enriched miRNA program associated with reduced lymphocyte-related signatures and increased suppressive myeloid features.

### DMG-associated miRNAs converge on multiple immune-regulatory pathways

Because cluster 6 was linked to both oncogenic miRNAs and suppressive TME features, we investigated whether its constituent miRNAs converge on coordinated immune pathways. Predicted targets were identified for 40 of the 46 cluster 6 miRNAs using miRTarBase (annotated miRNAs) or miRanda (novel miRNAs). Target sets for 27 miRNAs were significantly enriched for immune-related pathways (FDR < 0.05), including T-cell function for 13 miRNAs, myeloid-cell function for 22, antigen presentation or major histocompatibility complex-associated processes for three, and cytokine signaling for six (**Figure 1E; Table S5A**). These categories were not mutually exclusive, supporting the potential for individual miRNAs to influence multiple components of the immune microenvironment.

We next integrated predicted miRNA targets with matched bulk RNA-seq profiles to determine whether these computationally predicted relationships were reflected in patient tumors. Genes differentially expressed between DMG and non-tumor brain were identified using the same statistical framework applied to the miRNA analysis **(Figures S4A and S4B; Tables S6A and S6B**). We then assessed Pearson correlations between the expression of cluster 6 miRNAs and their predicted targets, with multiple-testing correction across all tested pairs.

To evaluate coordinated pathway-level regulation, we calculated expression scores for predicted target genes within each enriched immune pathway and related these scores to miRNA expression. This analysis identified 73 significant miRNA-pathway relationships, all of which were negative (adjusted p < = 0.05), consistent with the expected direction of miRNA-mediated repression. The associations involved five cluster 6 miRNAs: miR-130a-3p, with 25 pathway associations; miR-210-3p, with nine; miR-25-3p, with one; miR-502-3p, with seven; and miR-532-5p, with 31. MiR-210-3p was inversely associated with target programs related to myeloid-cell homeostasis, myeloid differentiation, and type I interferon responses (r range = - 0.95 to −0.75, adjusted p range = 1.5e-04 to 0.036; **Figure S5A**). MiR-532-5p was inversely associated with target programs involved in T-cell immunity and antigen processing and presentation (r = −0.84, adjusted p range = 5.1e-03; **Figure S5B**). Thus, a subset of DMG-enriched miRNAs was associated with coordinated changes across broad immune pathways rather than with isolated target genes alone.

Individual miRNA-target relationships further connected cluster 6 with T-cell activation, antigen presentation, and myeloid-cell regulation. Among the 27 cluster 6 miRNAs with immune-enriched target sets, 14 were predicted to regulate genes involved in T-cell biology (**Figure 1F**). MiR-320b and miR-320c were associated with 25 predicted target genes involved in T-cell activation (Pearson r < 0 or inverse differential expression pattern and FDR < 0.05). Although the direct effects of miR-320b and miR-320c on T-cell function remain to be established experimentally, these associations nominate the miR-320 family as a potential regulator of adaptive immune programs in DMG.

Four cluster 6 miRNAs were associated with antigen-presentation networks (**Figure S5C**). Among these, miR-148a-3p was inversely associated with four antigen-presentation genes, including *ARL8B* (r = −0.72, adjusted p = 0.02). Nineteen cluster 6 miRNAs were also linked to target programs involved in myeloid-cell differentiation and function (**Figure S5D**). MiR-21-5p and miR-3065-5p demonstrated the broadest associations, involving 16 and 14 myeloid-related genes, respectively. Collectively, these analyses identify a DMG-associated miRNA program linked to multiple interacting components of antitumor immunity, including T-cell activation, antigen presentation, interferon signaling, and myeloid-cell differentiation.

### MB contains a distinct and underannotated immunoregulatory miRNA landscape

To determine whether immune-associated miRNA programs extend beyond DMG, we next analyzed MB tumors. Differential-expression analysis comparing MB with tumor-adjacent or healthy pediatric brain identified 45 upregulated and 18 downregulated miRNAs (|log fold change| > 0, Wald test false discovery rate [FDR] < 0.05; **Figures 2A, 2B, S4C, and S4D; Tables S2C and S2D**). Unsupervised clustering separated MB from non-tumor brain and identified six distinct miRNA expression clusters (**Figure 2C**), demonstrating that MB harbors a miRNA landscape distinct from that of non-tumor tissue.

**Figure 2.**
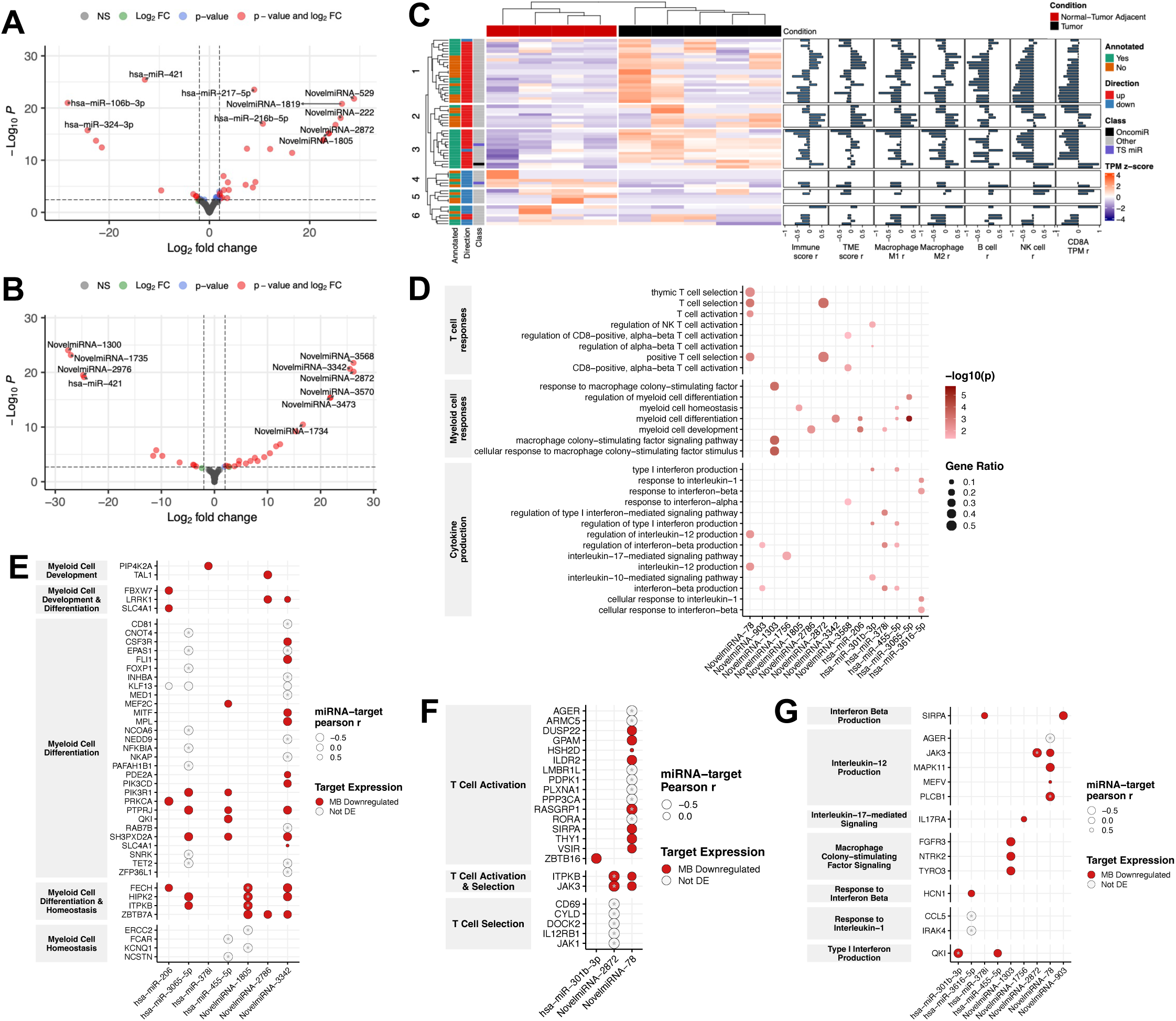
Medulloblastoma exhibits a distinct immune-associated miRNA landscape. (A-B) Volcano plots showing differential miRNA expression in medulloblastoma (MB) relative to healthy pediatric brain (**A**; n = 5/group) and tumor-adjacent brain (**B**; n = 3/group). Dashed lines indicate FDR < 0.05 and |log fold change| > 2. **(C)** Hierarchical clustering of differentially expressed miRNAs in MB and tumor-adjacent brain tissue. Adjacent bar plots depict correlations between miRNA clusters and inferred immune-cell abundance, immune scores, tumor microenvironment scores, and *CD8A* expression. **(D)** Gene Ontology enrichment analysis of predicted targets of MB cluster 1 miRNAs. **(E-G)** miRNA-target interaction networks associated with (**E**) myeloid-cell processes, (**F**) T-cell processes, and (**G**) additional immune-signaling pathways. Interactions were supported by inverse miRNA-target correlations and/or opposing differential expression patterns. Asterisks denote Pearson correlation *p* < 0.05.

A substantial proportion of the MB-associated miRNAs were previously unannotated. Of the 63 differentially expressed miRNAs, 26 lacked miRBase annotation (41.3%), compared with 29 of 170 miRNAs identified in DMG (17.1%; Fisher’s exact test, odds ratio [OR] = 3.4, 95% confidence interval [CI] = 1.7-6.8, p = 2.2e-04). This enrichment reveals a comparatively undercharacterized miRNA repertoire in MB and identifies candidate regulatory transcripts that would not be captured by analyses restricted to annotated miRNAs.

MB cluster 1 was most strongly associated with reduced antitumor immune signatures. This cluster contained 23 miRNAs, including 22 upregulated and one downregulated miRNA, and was associated with lower inferred lymphocyte and cytotoxic-cell abundance (CD8A expression mean r = −0.53, range = −0.96-0.03; **Figure 2C**). Twelve of the 23 cluster 1 miRNAs lacked miRBase annotation, and only three had previously been implicated in cancer. These findings identify cluster 1 as a largely unexplored MB-associated program linked to reduced immune activation.

We next examined whether cluster 1 miRNAs converge on immune-regulatory pathways. Predicted targets were identified for all 23 miRNAs, and targets of 15 miRNAs were significantly enriched for immune-related processes (FDR < 0.05; **Table S5B**). These included T-cell processes for four miRNAs, myeloid-cell processes for eight, and cytokine-related pathways for eight (**Figure 2D**). Integration with matched bulk RNA-seq identified inverse relationships with genes involved in myeloid-cell biology, T-cell function, and interferon and interleukin production or response (**Figures 2E–2G**).

MiR-3065-5p emerged as a candidate component of a shared myeloid-associated program. This miRNA was upregulated in both MB and DMG and was inversely associated with 14 genes involved in myeloid differentiation or function, including *TET2*, *FOXP1*, and *PIK3R1* (r range = - 0.96 to −0.94, adjusted p range = 1.7e-03 to 0.02; **Figure 2E**). Its recurrence across two histologies suggests that miR-3065-5p may mark a conserved myeloid-regulatory state across pediatric brain tumors, although functional perturbation will be required to establish direct regulation.

### Shared immune-associated miRNAs extend across pediatric CNS tumor histologies

To distinguish histology-specific dysregulation from regulatory programs shared across pediatric CNS tumors, we extended the analysis to EPN and LGG. Differential-expression analysis identified 31 upregulated and 19 downregulated miRNAs in EPN relative to adjacent or healthy brain tissue and 36 upregulated and 52 downregulated miRNAs in LGG relative to healthy brain (|log fold change| > 0, Wald test FDR < 0.05; **Figures S6A–S6C and S2E–S2F**).

Each histology retained a substantial tumor-specific component. Sixteen of the 31 miRNAs upregulated in EPN and 20 of the 36 upregulated in LGG were not upregulated in the other evaluated histologies (**Figure S6D**). These findings demonstrate that pediatric CNS tumors harbor distinct post-transcriptional programs that parallel their biological heterogeneity.

Despite this divergence, immune-associated networks were recurrent across tumor types. Targets of 22 EPN-upregulated and 24 LGG-upregulated miRNAs were enriched for immune-related Gene Ontology terms (FDR <0.05; **Figures S6E and S6F**). EPN shared six upregulated miRNAs with DMG cluster 6, whereas LGG shared nine upregulated miRNAs with MB cluster 1. Shared miRNAs included miR-21-5p, miR-210-3p, and miR-148a-3p, each of which was linked to immune-regulatory target programs in the preceding analyses. Thus, pediatric CNS tumors contain both histology-specific miRNA landscapes and recurrent miRNA programs associated with shared immune features of the TME.

### B7-H3 CAR T-cell therapy dynamically remodels miRNA and immune programs

Having identified immune-associated miRNA programs in patient tumors, we next asked whether these programs are dynamically remodeled during CAR T-cell therapy. We performed small RNA sequencing and bulk RNA sequencing on tumors from immunocompetent, high-grade glioma-bearing mice treated with B7-H3 CAR T cells containing a CD28 transmembrane and CD28 costimulatory domain, control CAR T cells containing a truncated endodomain, or no cellular therapy. Tumors were collected 14 and 21 days after treatment (**Figure S7; Table S7**). Group sizes were as follows: day 14, untreated n = 1, control CAR n = 3, and B7-H3 CAR n = 2; day 21, untreated n = 2, control CAR n = 3, and B7-H3 CAR n = 3.

PCA demonstrated treatment-associated separation in both miRNA and gene-expression profiles at days 14 and 21 (**Figures S8A and S8B**). Because the B7-H3 CAR T-cell groups contained two and three tumors at the respective time points, the longitudinal genomic analyses were considered exploratory and were used to nominate therapy-associated regulatory programs for subsequent validation.

To distinguish changes associated with therapeutic CAR activity from those associated with T-cell delivery or the control construct, we independently compared B7-H3 CAR-treated and control CAR-treated tumors with untreated tumors at each time point. After excluding miRNAs that also met differential-expression criteria in the control CAR comparison, 89 miRNAs remained significantly altered in B7-H3 CAR-treated tumors at one or both time points (|log fold change| > 0, Wald test FDR < 0.1; **Figures 3A and 3B; Table S8**). This filtering strategy identified changes preferentially associated with B7-H3 CAR T-cell treatment, although it did not constitute a formal treatment-by-time interaction test.

**Figure 3.**
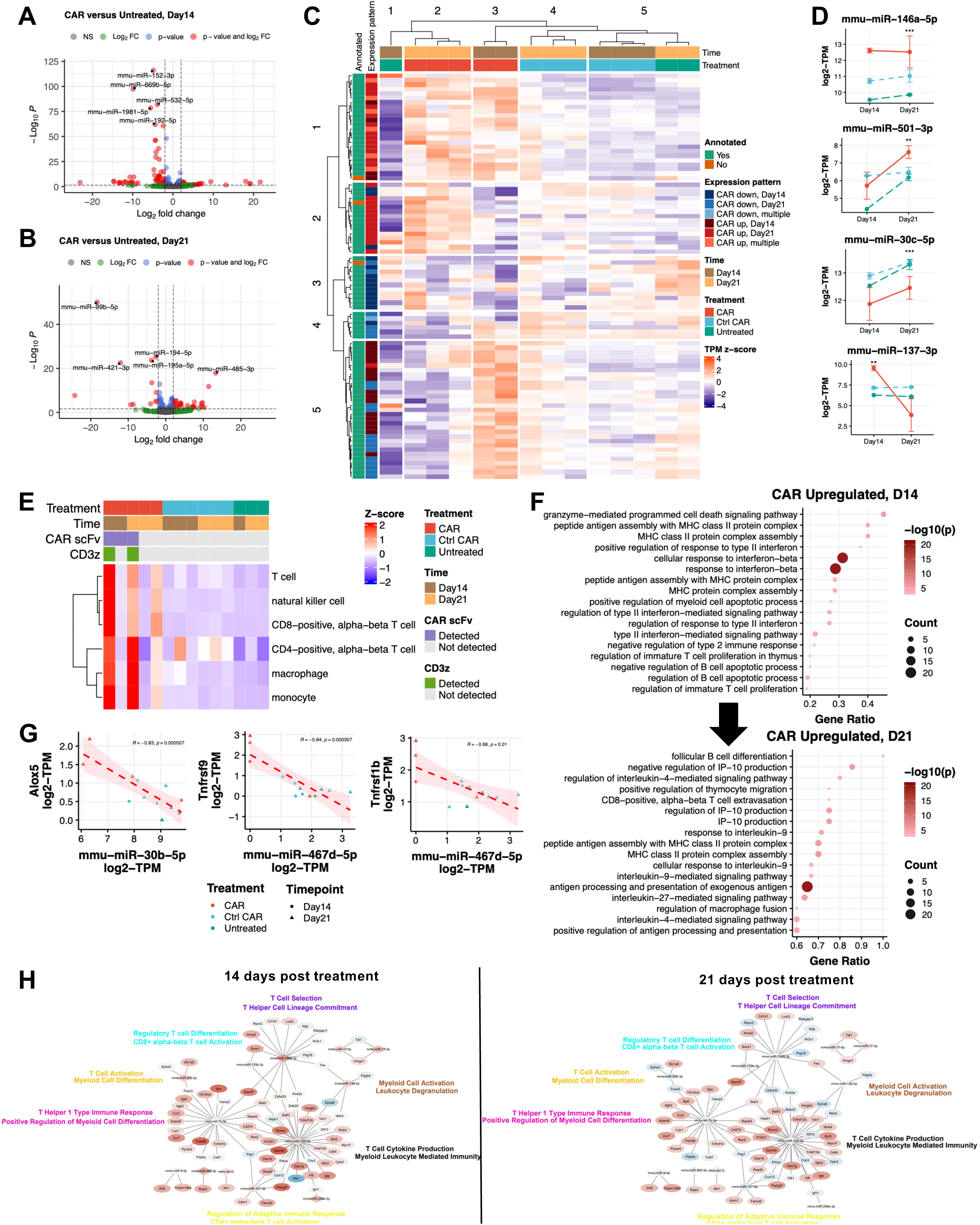
B7-H3 CAR T-cell therapy induces dynamic remodeling of miRNA and immune programs. (A-B) Volcano plots showing differential miRNA expression following B7-H3 CAR T-cell treatment relative to untreated tumors at day 14 (**A**) and day 21 (**B**). Day 14: n = 2 CAR-treated and n = 1 untreated tumors. Day 21: n = 3 CAR-treated and n = 2 untreated tumors. **(C)** Hierarchical clustering of therapy-associated differentially expressed miRNAs across all murine samples (n = 14). **(D)** Representative miRNAs illustrating the major temporal expression patterns observed following CAR T-cell therapy. Asterisks indicate adjusted *p* < 0.05. **(E)** Heatmap of inferred immune-cell populations from xCell2 deconvolution analysis together with detection of CAR transgene sequences in RNA-seq data. **(F)** Top immune-related Gene Ontology terms enriched among genes upregulated following B7-H3 CAR T-cell therapy at days 14 and 21. **(G)** Representative inverse correlations between therapy-regulated miRNAs and predicted immune-associated target genes, including miR-30b-5p:*Alox5*, miR-467d-5p:*Tnfrsf9*, and miR-467d-5p:*Tnfrsf1b*. **(H)** Integrated miRNA-gene interaction networks at early and late stages following CAR T-cell treatment. Triangles represent miRNAs and circles represent target genes. Colors correspond to log fold-change values (red, upregulated; blue, downregulated).

Unsupervised clustering revealed five temporal patterns of miRNA expression following therapy (**Figures 3C and 3D; Table S9**). These included miRNAs that were broadly 1) upregulated in B7H3 treatment group at both time points (cluster 1, N = 24); 2) upregulated in B7H3 at Day21 (cluster 2, N = 16); 3) downregulated in B7H3 at both time points (cluster 3, N = 12); or 4) upregulated in B7H3 at day 14 (clusters 4 and 5, N = 37). The distinct trajectories indicate that CAR T-cell therapy induces both sustained and time-restricted changes in the tumor miRNA landscape.

Parallel transcriptomic analysis revealed substantial temporal remodeling of immune programs. At day 14, 46 genes were uniquely upregulated and 291 downregulated in B7-H3 CAR-treated tumors relative to untreated tumors after removal of changes detected in the control CAR comparison. By day 21, 626 genes were upregulated and 1,063 downregulated (**Figures S8C and S8D**). Immune deconvolution identified higher inferred immune-cell content in one of two B7-H3 CAR-treated tumors at day 14 and two of three at day 21, whereas untreated and control CAR-treated tumors generally had low inferred immune abundance (**Figure 3E**). CAR transcripts were inconsistently detected and did not correspond to the magnitude of inferred immune infiltration. Given the cohort size, these observations illustrate heterogeneity among treated tumors but do not establish a group-level difference in immune-cell abundance.

Pathway analysis nevertheless identified biologically distinct phases of treatment-associated transcriptional remodeling (**Table S10**). Genes upregulated at day 14 were enriched for T-cell proliferation, antigen presentation, and interferon signaling (Fisher’s exact test [FET] OR > 1, FDR < 0.05) consistent with an early effector-associated response. Genes upregulated at day 21 were enriched for T-cell recruitment and interleukin signaling (FET OR > 1, FDR < 0.05), suggesting a later state characterized by immune trafficking and inflammatory remodeling (**Figure 3F**). Together, these results connect temporal changes in miRNA expression with early effector and later recruitment-associated transcriptional states following B7-H3 CAR T-cell therapy.

### Therapy-responsive miRNAs are associated with temporally distinct immune programs

We next integrated therapy-responsive miRNAs with immune-related genes to identify candidate regulators of the temporal transcriptional response. We prioritized predicted miRNA-target pairs in which the miRNA and target gene changed in opposite directions and their expression was significantly inversely correlated (**Figure 3G**).

MiR-467d-5p emerged as a candidate regulator of the early immune response. MiR-467d-5p was significantly downregulated at day 14 (log fold change = −10.11, FDR = 2.6e-04) and was inversely associated with concurrently upregulated immune-related genes (r range = −0.84-- 0.59, adjusted p range = 3.10e-04-0.03; **Figure S9**). Predicted targets included *Tnfrsf9*, which regulates T-cell activation and persistence; *Tnfrsf1b*, a mediator of TNF signaling; the interferon-responsive genes *Oas3*, *Apobec3*, and *Ifi213*; *Clec7a*, associated with activated myeloid states; and genes involved in cellular proliferation and tissue remodeling, including *Cdc6*, *Loxl3*, and *Ddx21*. These relationships suggest that reduced miR-467d-5p expression accompanies activation of multiple immune and inflammatory pathways early after therapy.

A distinct candidate emerged during the later response. MiR-30b-5p was significantly downregulated at day 21 (log fold change = −2.34, FDR = 1.97e-04) and inversely associated with increased *Alox5* expression (r = −0.83, adjusted p = 5.1e-04). *Alox5* encodes a mediator of leukotriene synthesis linked to inflammatory signaling and immune-cell recruitment. These data nominate miR-467d-5p and miR-30b-5p as candidate regulators of temporally distinct immune programs, but direct regulation will require functional validation.

### Human DMG-associated miRNAs are recapitulated during CAR T-cell therapy

We next asked whether miRNA programs identified in human tumors are conserved during therapeutic remodeling in immunocompetent models. Of 141 annotated miRNAs differentially expressed in human DMG, 63 had annotated murine orthologs and 61 of these were detected in the murine dataset (**Figure 4A**). Twenty orthologous miRNAs were differentially expressed following B7-H3 CAR T-cell therapy at one or both time points, representing significant enrichment relative to the tested miRNA background (Fisher’s exact test, OR = 3.41, 95% CI = 1.78–6.40, p = 1.45 × 10□□).

**Figure 4.**
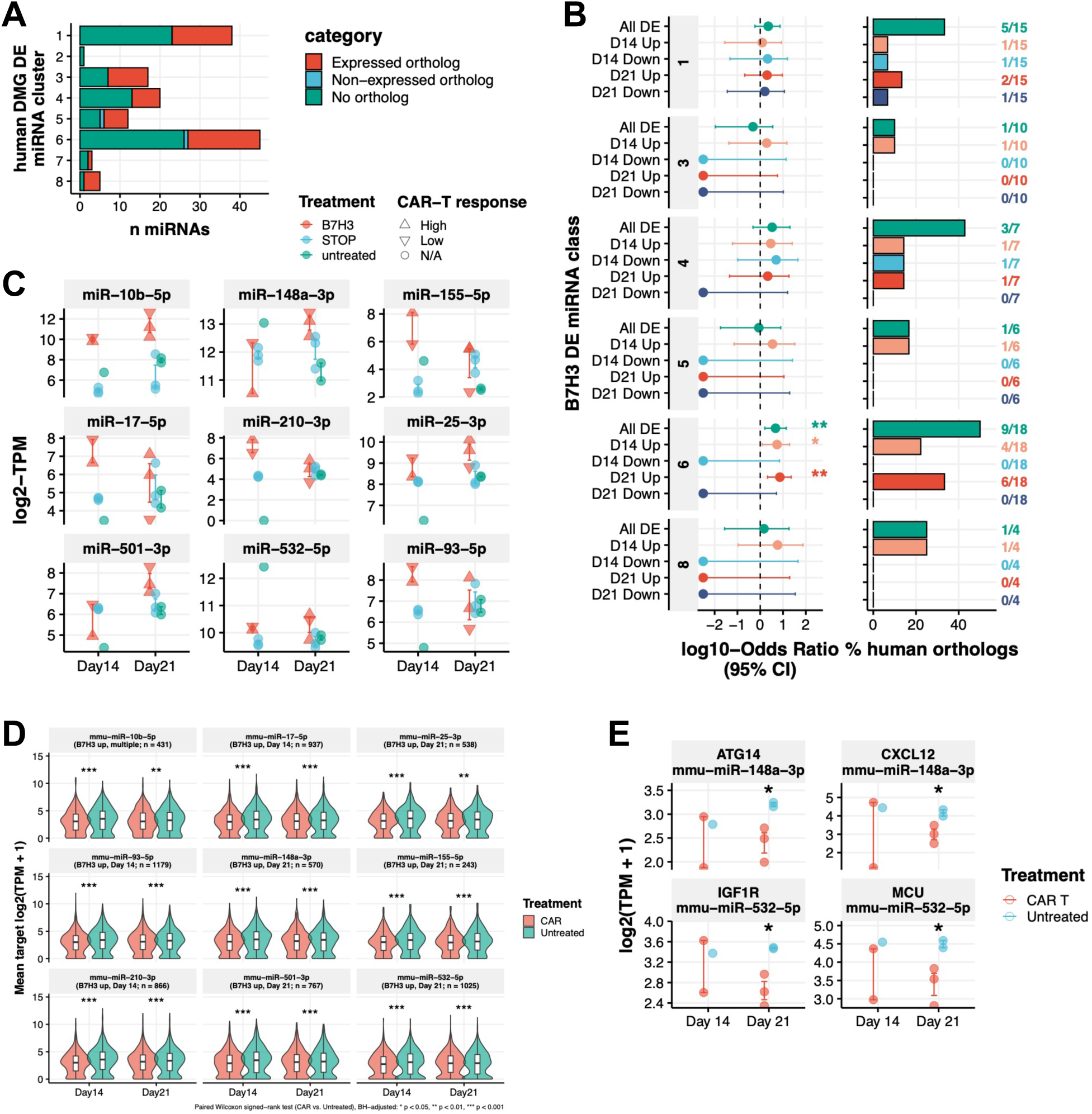
Human tumor-associated miRNA programs are recapitulated during CAR T-cell therapy. **(A)** Number of differentially expressed human DMG miRNAs with detectable murine orthologs, stratified by human miRNA cluster assignment. **(B)** Enrichment of human DMG-derived orthologous miRNAs among CAR T-responsive murine miRNAs. Bars indicate the percentage of orthologous miRNAs within each expression category. *p* < 0.05; **p** < 0.01. **(C)** Expression of CAR T-responsive cluster 6 orthologous miRNAs across treatment groups and time points. **(D)** Distribution of predicted target-gene expression in CAR T-treated and untreated tumors. Statistical significance was evaluated using Wilcoxon rank-sum testing. **(E)** Expression of representative immune-associated targets of miR-148a-3p (*Atg14* and *Cxcl12*) and miR-532-5p (*Igf1r* and *Mcu*) following CAR T-cell therapy. Asterisks indicate Wilcoxon rank-sum *p* < 0.05.

Cluster 6 was the only human DMG cluster significantly enriched for therapy-responsive orthologs (OR = 4.7, 95% CI = 1.6-13.8, adjusted p = 2.1e-03; **Figure 4B**). Nine of the 18 orthologous cluster 6 miRNAs were upregulated after treatment at day 14, day 21, or both. Because cluster 6 was associated with reduced lymphocyte-related signatures and increased suppressive myeloid features in human DMG, its induction after CAR T-cell therapy raises the possibility that components of this program are engaged as part of an adaptive response to immune pressure.

We assessed whether induction of these nine miRNAs was accompanied by changes in their predicted targets. Predicted target sets for each miRNA shifted toward lower expression following B7-H3 CAR T-cell therapy at both time points (paired Wilcoxon rank-sum test FDR < 4.0 × 10 ³). Across the combined set of 4,895 predicted targets, target genes were significantly enriched among genes downregulated after therapy at Day 21 (Fisher’s exact test, OR = 2.20, 95% CI = 1.86-2.60, FDR = 4.9e-18; **Figure S10A**). These results are consistent with aggregate repression of predicted targets but do not establish direct targeting by individual miRNAs.

Immune-annotated targets demonstrated a more complex response. Among 871 immune-related targets, only those of miR-25-3p, miR-210-3p, and miR-501-3p were significantly downregulated following therapy at Day 14 (**Figure S10B**). However, these genes were enriched among downregulated genes following therapy at Day 21 (N = 42, OR = 2.23, 95% CI = 1.78-3.56), providing evidence for miRNA-mediated repression despite broad transcriptional activation of immune pathways. Immune-pathway targets of three individual miRNAs remained enriched among genes downregulated after therapy. These included four miR-155-5p targets (OR = 5.56, 95% CI = 1.86-13.90, FDR = 4.0e-03), seven miR-148a-3p targets (OR = 2.81, 95% CI = 1.14–6.02, FDR = 0.086), and nine miR-93-5p targets (OR = 2.17, 95% CI = 1.06-4.10, FDR = 0.095).

Among miR-148a-3p targets, *Atg14* and *Cxcl12* were downregulated following therapy (**Figure 4E**; log fold changes = −0.87 and −1.24, FDR = 0.030 and 0.027, respectively). MiR-532-5p was also predicted to target *Igf1r* and *Mcu*, which participate in T-cell metabolic and mitochondrial signaling. Collectively, these analyses demonstrate that a human DMG-associated miRNA program is dynamically engaged during CAR T-cell therapy and identify candidate regulatory relationships that may influence immune-cell recruitment, metabolic fitness, and effector function.

### Modulation of miR-149 improves CAR T-cell fitness during repeated antigen exposure

To determine whether candidate miRNAs identified through our integrated tumor and therapy-response analyses could be therapeutically leveraged to enhance CAR T-cell function, we selected miR-149 for functional validation. MiR-149 was prioritized because it is predicted to regulate multiple pathways implicated in T-cell dysfunction and exhaustion, including PD-1, TIM-3, BTLA, and *Foxp1*. We generated murine B7-H3 CAR T cells co-expressing a miR-149 overexpression cassette (miR-149-B7-H3 CAR) and compared them with matched parental B7-H3 CAR T cells lacking the miRNA cassette (**Figure 5A**).

**Figure 5.**
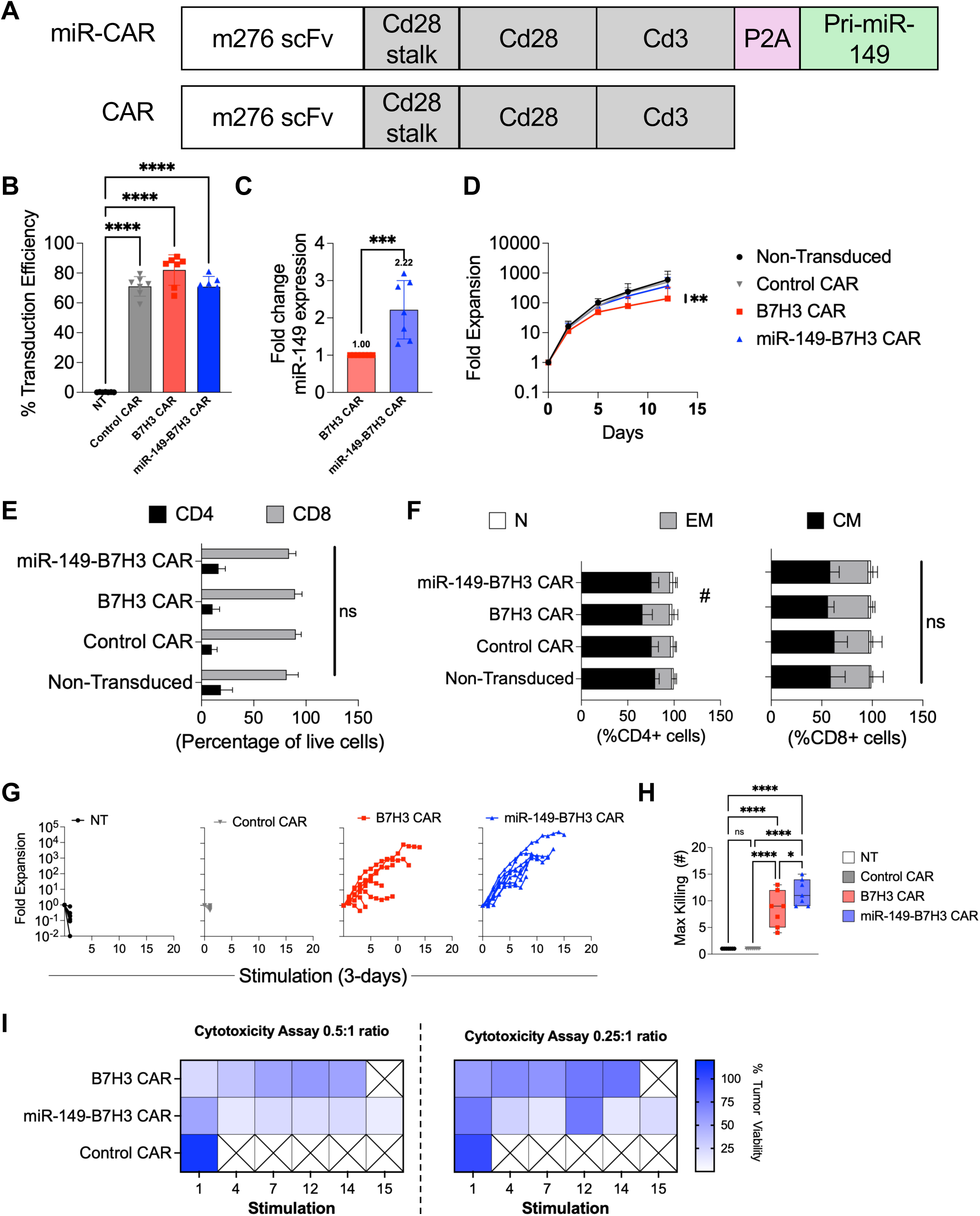
miR-149 enhances CAR T-cell fitness during chronic antigen exposure. **(A)** Schematic of parental B7-H3 CAR and miR-149-B7-H3 CAR constructs. The miR-149-B7-H3 CAR contains dual miR-149-5p and miR-149-3p expression cassettes driven by the U6 and H1 promoters, respectively. **(B)** Transduction efficiency of non-transduced (NT), control CAR, B7-H3 CAR, and miR-149-B7-H3 CAR T cells. **(C)** Relative miR-149 expression measured by droplet digital PCR in parental and miR-149-engineered CAR T cells. **(D)** Expansion during standard in vitro manufacturing conditions. **(E)** CD4□ and CD8□ T-cell composition of generated products. **(F)** Baseline distribution of naïve (N), central memory (CM), and effector memory (EM) T-cell subsets among CD4 and CD8 populations. **(G)** Representative serial tumor-rechallenge assay showing cumulative expansion of T cells subjected to repeated antigen stimulation every 3 days. **(H)** Maximum sequential killing capacity achieved during serial restimulation assays. Each point represents an independent donor. **(I)** Cytotoxic activity during prolonged antigen exposure at effector-to-target ratios of 0.5:1 and 0.25:1. Heatmaps depict residual tumor viability across sequential rounds of stimulation. Data are presented as mean ± SD unless otherwise indicated. Statistical analyses are described in STAR Methods. ns, not significant. #*p* < 0.05; *p* < 0.05; **p** < 0.01; ***p*** < 0.001; **p** < 0.0001.

MiR-149 overexpression did not adversely affect CAR T-cell manufacturing or baseline phenotype. Both constructs achieved efficient transduction, and incorporation of the miR-149 cassette resulted in a significant increase in miR-149 expression (**Figure 5C**; mean fold-change = 2.22 ± 0.78, *p* = 0.0003) without altering CAR expression levels (**Figure 5B**). Likewise, miR-149 overexpression did not affect cell expansion during manufacturing (**Figure 5D**), CD4:CD8 composition (**Figure 5E**), or the distribution of naïve, central memory, and effector memory T-cell subsets at baseline (**Figure 5F**; *p* > 0.05 for all comparisons except CD4 central memory cells, NT versus B7-H3 CAR, *p* = 0.0019, and B7-H3 CAR versus miR-149-B7-H3 CAR, *p* = 0.0429). These findings indicate that miR-149 does not substantially alter CAR T-cell differentiation during production.

We next asked whether miR-149 could improve CAR T-cell performance under chronic antigen exposure, a major driver of dysfunction in solid tumors. Using a serial tumor restimulation assay, CAR T cells were challenged with fresh tumor targets every three days until loss of proliferative capacity or cytotoxic activity. While non-transduced and control CAR T cells rapidly lost expansion potential and antitumor activity, miR-149-B7-H3 CAR T cells demonstrated greater cumulative expansion and sustained proliferative capacity across repeated stimulations compared with parental B7-H3 CAR T cells (**Figure 5G**).

This enhanced fitness translated into prolonged functional activity. MiR-149-B7-H3 CAR T cells remained effective through significantly more rounds of sequential tumor challenge, achieving a mean of 12 productive killing cycles compared with 8 cycles for parental B7-H3 CAR T cells (**Figure 5H**; B7-H3 CAR versus miR-149-B7-H3 CAR, *p* = 0.04). As expected, both CAR T-cell products significantly outperformed non-transduced and control CAR T cells (*p* < 0.0001 for all comparisons), whereas no difference was observed between non-transduced and control CAR groups (*p* > 0.9).

Although miR-149-B7-H3 CAR T cells exhibited slightly slower tumor killing during the initial stimulation, they maintained cytotoxic activity throughout prolonged antigen exposure, whereas conventional B7-H3 CAR T cells progressively lost function, particularly at lower effector-to-target ratios (**Figure 5I**). These findings suggest that miR-149 promotes a more durable functional state characterized by sustained proliferation and preserved antitumor activity under conditions of chronic stimulation.

Collectively, these data provide functional validation of a miRNA identified through integrated analyses of pediatric brain tumors and treatment-associated immune remodeling. More broadly, they demonstrate that dysregulated miRNA programs can be therapeutically harnessed to improve engineered T-cell fitness. The ability of miR-149 to enhance CAR T-cell durability supports the concept that modulation of regulatory miRNA networks may represent a complementary strategy to conventional receptor-or pathway-specific approaches for overcoming T-cell dysfunction in pediatric brain tumors.

## Discussion

Pediatric brain tumors encompass biologically diverse diseases with equally heterogeneous immune microenvironments (7,8,17). Recent large-scale immune profiling has shown that immune activation varies both across and within tumor types, supporting evaluation of the tumor immune microenvironment as a biological feature distinct from histologic or molecular diagnosis (17). However, the regulatory mechanisms that establish these immune states and change in response to therapy remain incompletely defined. Here, we identify immunomodulatory microRNA programs associated with pediatric brain tumor subtype, immune microenvironment composition, and therapeutic response. By integrating miRNA and transcriptomic data with functional studies, our findings extend descriptive immune profiling to reveal candidate regulatory networks that can be experimentally manipulated to improve antitumor T-cell function.

A central finding of this study is that pediatric brain tumors exhibit distinct patterns of immunomodulatory miRNA expression relative to non-tumor brain tissues, with both shared and tumor-specific changes across diffuse midline glioma, medulloblastoma, high-grade glioma, and other pediatric CNS tumor subtypes. This heterogeneity is consistent with evidence that pediatric brain tumors cannot be uniformly classified as immunologically “cold” and that immune activation can vary substantially even within a molecularly defined tumor entity (17). Our data add another regulatory dimension to this emerging framework. Rather than measuring immune composition or inflammatory gene expression alone, miRNA profiling identifies upstream post-transcriptional programs capable of coordinating multiple immune and tumor-associated pathways. Thus, pediatric brain tumor immune states may be distinguished not only by the abundance or activation state of immune cells but also by the regulatory networks that sustain those states.

This distinction is important because most approaches to overcoming tumor-associated immune suppression focus on a single receptor, cytokine, signaling pathway, or cellular population (9,10,18). Such strategies may be insufficient in pediatric brain tumors, where immune dysfunction arises from interconnected tumor-intrinsic, myeloid, and lymphoid programs (17). miRNAs are well suited to addressing this complexity because individual miRNAs can regulate multiple transcripts within related biological pathways. Our integrated miRNA–mRNA analyses therefore provide more than a catalog of differentially expressed miRNAs: they generate testable regulatory networks linking miRNA dysregulation to immune recruitment, activation, differentiation, and suppression. The conceptual innovation of this work is the use of miRNAs as systems-level regulators of the tumor immune microenvironment rather than as isolated diagnostic or prognostic markers.

Our analysis also demonstrates that miRNA programs are dynamic and change following CAR T-cell therapy. Profiling treated tumors identified therapy-associated miRNAs linked to myeloid and T-cell programs, indicating that immune-directed therapy can modify the regulatory architecture of the tumor microenvironment. This finding is significant because most pediatric brain tumor miRNA studies have focused on differences between tumor subtypes or between tumor and normal tissue (19–21). By integrating baseline human tumor profiles with therapy-response models, our study distinguishes miRNAs associated with tumor identity from those associated with immune remodeling after treatment. This framework may help identify biomarkers that reflect whether a therapy has engaged the tumor microenvironment, even when changes in tumor burden alone provide an incomplete measure of biological response.

The functional evaluation of miR-149 provides proof of concept that candidates emerging from these integrated analyses can be therapeutically leveraged. Previous work in a breast cancer model showed that miR-149-3p was predicted to target the 3′ untranslated regions of transcripts encoding PD-1, TIM-3, BTLA, and FOXP1 (22). Delivery of a miR-149-3p mimic reduced expression of these inhibitory regulators while increasing T-cell proliferation, effector cytokine secretion, and target-cell killing (22). Our findings extend this prior observation into engineered CAR T cells and a pediatric brain tumor context. Instead of applying a transient miRNA mimic to conventional CD8+ T cells, we incorporated a miR-149 cassette into a B7-H3 CAR platform and evaluated function during repeated tumor exposure. miR-149 expression did not compromise transduction, cellular composition, or overall baseline differentiation, supporting the feasibility of integrating a regulatory miRNA into CAR T-cell manufacturing.

The principal benefit of miR-149 emerged during chronic rather than initial antigen exposure. Although miR-149-modified CAR T cells did not demonstrate uniformly greater short-term activity, they maintained expansion and tumor killing through more rounds of serial stimulation than parental B7-H3 CAR T cells. This distinction is biologically meaningful because persistent antigen exposure can progressively reduce T-cell proliferation, cytokine production, and cytotoxic function while increasing the expression of multiple inhibitory receptors. Solid tumor microenvironments further limit CAR T-cell persistence and activity through combined antigenic, trafficking, metabolic, and immunosuppressive barriers. Our data suggest that miR-149 does not simply increase immediate CAR T-cell potency. Rather, miR-149 supports a more durable functional state that becomes apparent under repeated antigenic challenge, a condition more relevant to the sustained pressure encountered by CAR T cells in solid tumors.

This finding highlights a broader innovation in CAR T-cell design. Many current engineering strategies address one inhibitory mechanism at a time, including checkpoint disruption, cytokine armoring, dominant-negative receptors, or modification of individual transcriptional and epigenetic regulators (10). In contrast, miRNA engineering offers a compact means of coordinately modulating several functionally connected targets. The sustained activity of miR-149-B7-H3 CAR T cells is consistent with this network-level model, although the contribution of each predicted target remains to be established. The current results should therefore not be interpreted as demonstrating that PD-1, TIM-3, BTLA, or FOXP1 individually mediates the observed phenotype. Rather, the data support the premise that miR-149 can improve functional durability during chronic antigen exposure, potentially through coordinated regulation of multiple pathways that govern T-cell fitness.

The study adds three advances to the field. First, it provides an integrated view of immunomodulatory miRNA dysregulation across multiple pediatric brain tumor types, linking miRNA expression to features of the immune microenvironment rather than examining miRNAs solely as tumor-intrinsic biomarkers. Second, it incorporates treatment-associated profiling to identify miRNAs that change as the tumor microenvironment responds to CAR T-cell therapy. Third, it connects discovery to therapeutic application by demonstrating that one prioritized miRNA can be incorporated into a CAR construct to improve functional durability during serial tumor challenge. Together, these advances establish a discovery-to-engineering framework in which miRNA profiles are used to understand tumor immune biology, monitor therapy-associated remodeling, and identify candidates for improving cellular immunotherapy.

Several limitations warrant consideration. The functional studies focused on miR-149 and therefore do not establish whether other candidates identified by the integrated analyses will produce similar effects. The serial restimulation assay models repeated antigen exposure but does not reproduce the full cellular, spatial, and metabolic complexity of the brain tumor microenvironment. In addition, although previous work supports regulation of PD-1, TIM-3, BTLA, and FOXP1 by miR-149-3p, the specific mature miR-149 strand expressed from our construct and the direct molecular targets responsible for enhanced CAR T-cell durability require experimental confirmation. Future work should combine target-validation studies with transcriptomic and functional profiling after repeated stimulation and determine whether the in vitro durability phenotype translates into improved persistence and tumor control in vivo.

In summary, our study identifies miRNA regulation as an important and therapeutically actionable layer of pediatric brain tumor immunity. The findings move beyond describing which miRNAs are altered by defining how miRNA programs relate to immune context, change following therapy, and can be harnessed to improve engineered T-cell function. Most importantly, miR-149 engineering enhanced CAR T-cell durability under repeated antigen exposure without substantially disrupting baseline product characteristics. These results support a shift from modifying single suppressive pathways toward leveraging miRNA networks to coordinate multiple determinants of immune dysfunction. This framework provides a foundation for developing miRNA-guided biomarkers and next-generation CAR T-cell therapies designed to remain functional within the complex and evolving microenvironments of pediatric brain tumors.

## Materials and Methods

### Human pediatric CNS tumor specimens

Human tissue specimens were obtained from the Children’s National Hospital Brain Tumor Repository under an Institutional Review Board-approved protocol (IRB #1339). All specimens were collected postmortem, flash frozen, and stored until use.

The cohort included 27 pediatric CNS tumor specimens, 18 tumor-adjacent non-neoplastic brain specimens, and 5 non-tumor pediatric brain specimens. Tumor-adjacent specimens were obtained from anatomically distinct regions without histopathologic evidence of tumor. Non-tumor controls were obtained from pediatric donors without brain cancer. Tumor histologies included diffuse midline glioma (DMG), medulloblastoma, ependymoma, high-grade glioma, low-grade glioma, and ATRT. Clinical and molecular characteristics are provided in **Table S1**.

Approximately 3 mg of tissue was processed from each specimen. Total RNA, including small RNAs, was isolated using the miRNeasy Tissue/Cells Advanced Micro Kit (QIAGEN, Cat# 217084). RNA concentration and purity were assessed using a NanoDrop 2000 spectrophotometer (Thermo Fisher Scientific, Cat# ND-2000).

### Murine glioma model and CAR T-cell treatment

Animal studies were approved by the Children’s National Hospital Institutional Animal Care and Use Committee (IACUC protocol #30512).

Female and male C57BL/6J mice (Jackson Laboratory, Stock No. 000664), 6–8 weeks of age, were implanted intracranially with 1 × 10^5 GL261 murine glioma cells (DSMZ German Collection Laboratory; ACC 802) using stereotactic guidance. Cells were suspended in phosphate-buffered saline and injected into the cerebral cortex of syngeneic mice as described previously (18).

Ten days after implantation, mice were randomized to receive no treatment, B7-H3 CAR T cells, or control CAR T cells. A total of 2 × 10^6 CAR T cells were administered intracranially. Control CAR T cells recognized B7-H3 but lacked functional downstream signaling due to a premature stop codon within the intracellular signaling domain.

Brains were harvested 4, 11, or 18 days after treatment (days 14, 21, and 28 after implantation, respectively). Tumor-bearing cortical tissue was isolated and processed for RNA extraction using the miRNeasy Micro Kit (QIAGEN, Cat# 217084).

### Generation of murine CAR T cells

CD3+ T cells were isolated from murine splenocytes using the Pan T Cell Isolation Kit II (Miltenyi Biotec, Cat# 130-095-130). Following ACK red blood cell lysis (Quality Biological, Cat# 118-156-101), T cells were activated using plate-bound anti-CD3 (BD Biosciences, Cat# 553057) and anti-CD28 (BD Biosciences, Cat# 553294) antibodies in complete RPMI supplemented with 50 U/mL human IL-2 (NIH/NCI, Ro 23-6019).

Two days after activation, T cells were transduced on RetroNectin-coated plates (Takara Bio, Cat# T100B) with retroviral vectors encoding either the parental B7-H3 CAR or a modified B7-H3 CAR construct co-expressing miR-149.

The parental construct contained the m276 anti-B7-H3 scFv, CD28 hinge/transmembrane and costimulatory domains, and a CD3ζ signaling domain (9). The miR-149 construct contained the B7-H3 CAR cassette with dual miR-149-5p and miR-149-3p expression cassettes driven by U6 and H1 promoters, respectively.

Transduced cells were expanded in complete RPMI supplemented with IL-2 and used for downstream analyses between days 4 and 6 after transduction.

### CAR T-cell phenotyping and transduction efficiency

CAR expression was detected using FITC-conjugated anti-human IgG F(ab’) fragment-specific antibody (Jackson ImmunoResearch, Cat# 109-096-006).

T-cell phenotype was assessed by flow cytometry using antibodies against CD4 (BD Biosciences, Cat# 552051), CD8 (BioLegend, Cat# 100742; BD Biosciences, Cat# 553030), CD62L (BD Biosciences, Cat# 560513), CD44 (BD Biosciences, Cat# 553134), PD-1 (BioLegend, Cat# 135220), TIM-3 (BioLegend, Cat# 134019), and TIGIT (BioLegend, Cat# 142106). Cell viability was assessed using LIVE/DEAD Aqua (Invitrogen, Cat# L34966).

Samples were acquired using a CytoFLEX flow cytometer (Beckman Coulter) and analyzed using FlowJo (version 10).

MiR-149 expression was quantified by droplet digital PCR using Advanced mmu-miR-149-5p and mmu-miR-149-3p Taqman Assays (Cat #A25576) on a QX 200 Droplet Reader.

### Repeat stimulation assay

To assess CAR T-cell durability under chronic antigen exposure, serial tumor restimulation assays were performed. Briefly, 5 × 10^5 GL261 tumor cells were plated in 24-well plates in 1 mL culture medium and allowed to adhere for 2–6 h at 37°C.

CAR T cells were washed once with cytokine-free medium, and 1 × 10^6 T cells were added to each well at an initial effector-to-target ratio of 2:1.

After 3–4 days, T cells were harvested by gentle pipetting and quantified using a hemocytometer. Fresh GL261 targets were replated, and recovered T cells were transferred at the same effector-to-target ratio. This process was repeated every 3–4 days until measurable proliferation and/or tumor killing could no longer be detected.

When fewer than 1 × 10^6 T cells remained but antitumor activity persisted, all recovered cells were replated at the maximum achievable effector-to-target ratio. The maximum sequential killing capacity was defined as the total number of productive stimulation cycles completed before loss of measurable antitumor activity.

### Cytotoxicity assay

Tumor cell killing was quantified using the CellTiter 96 AQueous One Solution MTS assay (Promega, Cat# G3581). GL261 cells were plated in tissue-culture-treated 96-well plates and allowed to adhere for 2–6 h before addition of T cells.

CAR T cells were added at effector-to-target ratios of 2:1, 1:1, 0.5:1, 0.25:1, and 0.125:1. Tumor-only and media-only wells served as controls. All conditions were performed in technical triplicate.

Following 3–4 days of co-culture, non-adherent T cells were removed and MTS reagent was added according to the manufacturer’s instructions. Plates were incubated for 2 h at 37°C, and absorbance was measured at 492 nm using a SPARK plate reader (Tecan).

Tumor viability was calculated as:

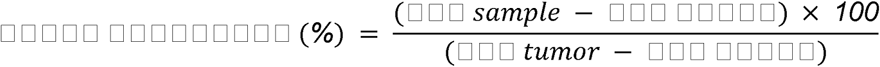

### Small RNA sequencing

Small RNA library preparation and sequencing were performed by Azenta Life Sciences using the NEBNext Multiplex Small RNA Library Prep Set for Illumina.

Libraries were sequenced on an Illumina NovaSeq X Plus instrument at a mean depth of 27 million reads per sample (range 13–140 million reads).

Reads were processed using Trimmomatic v0.30. Adapter sequences, bases with quality scores below 20, and reads shorter than 18 bp were removed. Filtered reads were aligned to the Rfam database (v15.0) using BLASTN (v2.16.0). Novel miRNAs were identified using miRDeep2 v2.0.0.7 following alignment to GRCh38 or GRCm39. Known miRNAs were annotated using miRBase (Release 22).

### Bulk RNA sequencing

Libraries were prepared by Azenta Life Sciences using the NEBNext Ultra II RNA Library Prep Kit and sequenced on an Illumina NovaSeq platform using paired-end 150-bp sequencing.

Human samples were sequenced to a mean depth of 35 million reads per sample (range 20–48 million reads). Murine samples were sequenced to a mean depth of 19 million reads per sample (range 15–25 million reads).

Reads were processed using the Kids First DRC RNA-seq workflow v4. Reads were trimmed using Cutadapt v3.4, aligned using STAR v2.7.10a, and quantified using RSEM v1.3.1 with GENCODE v38 (human) or GENCODE vM39 (mouse) annotations. Murine analyses used a customized GRCm39 reference genome containing B7-H3 CAR and control CAR transgene sequences.

### Differential expression analysis

Differential expression analyses were performed using DESeq2 (v1.48.2) in R (v4.5.0).

For human analyses, histology-specific negative binomial generalized linear models were fitted separately for each tumor type. Paired tumor-adjacent analyses included patient identity as a covariate. Genes and miRNAs with an FDR < 0.05 at any fold change value were considered differentially expressed.

For murine analyses, models were fitted separately at days 14 and 21. Features were classified as CAR-associated if they met an FDR < 0.10 for B7-H3 CAR versus untreated and an FDR > 0.10 for control CAR versus untreated.

### Immune deconvolution

Human samples were analyzed using immunedeconv (**v2.1.3**) with xCell and quanTIseq. Murine samples were analyzed using xCell2 (**v1.5.0**) with the ImmGen reference dataset.

Pearson correlation analyses were performed between miRNA expression and inferred immune-cell abundance, immune scores, and immune-associated gene-expression signatures.

### miRNA target prediction and network analysis

Experimentally validated miRNA targets were obtained from miRTarBase (v10.0). Targets of novel miRNAs were predicted using miRanda v3.3a (score >145; energy <−20 kcal/mol).

Gene Ontology enrichment analyses were performed using topGO (v2.60.1) with Benjamini-Hochberg correction. Terms with an FDR < 0.05 were considered significant.

miRNA-target relationships were evaluated using Pearson correlation analyses. Networks were visualized using Cytoscape 3.10.4, ClueGO 2.5.10, and CluePedia 1.5.10.

### Human-mouse miRNA orthology analysis

Human DMG-associated miRNAs were mapped to murine orthologs using miRBase nomenclature (Release 22). Ortholog enrichment among CAR-responsive murine miRNAs was assessed using two-sided Fisher’s exact tests.

### Statistical analysis

Statistical analyses for Figure 5 experiments were performed using GraphPad Prism (version 11.0.2). Differential expression analyses utilized Wald tests with Benjamini-Hochberg correction. Gene Ontology enrichment analyses were performed using Fisher’s exact tests. Pearson correlation was used for miRNA-target and immune-association analyses. Statistical tests, sample sizes, and significance thresholds are provided in the corresponding figure legends. Unless otherwise stated, adjusted p values < 0.05 were considered statistically significant. All R and package versions can be found in the Dockerfile associated with the GitHub repository: https://github.com/rokitalab/PedBrainTumor-miR-TME/blob/main/Dockerfile.

## Data availability

Raw data are undergoing processing for deposition into dbGAP. Processed data are available through data download script from the GitHub repository below.

## Code availability

All original code and manuscript figures and tables are available in the GitHub repository (https://github.com/rokitalab/PedBrainTumor-miR-TME) and Zenodo repository (DOI 10.5281/zenodo.23397057)(23).

## Acknowledgements

We are deeply grateful to the patients and families whose generous donation of tumor and brain tissue made this study possible. We thank the Children’s National Hospital Brain Tumor Biorepository, the Children’s Brain Tumor Network, The Swifty Foundation, and Gift from a Child who, together, enabled specimen collection, processing, and distribution.

We also thank the anonymous philanthropic donors to the Children’s National Hospital Brain Tumor Institute for supporting the bioinformatics analyses in this study, and Dr. Brian Rood and Dr. Javad Nazarian for their scientific support and contributions to the biorepository and research infrastructure that enabled this work.

## Author Contributions

D.H., K.C., and R.J.C. conceived the study. D.H., R.J.C., B.S., K.C., A.S., and A.B. developed methodology. D.H., K.C., R.J.C., B.S., A.S., and A.B. performed data analysis. K.C., R.J.C. and B.S. conducted experiments. D.H. and J.L.R. provided resources and supervision. K.C., R.J.C., B.S., and J.L.R. curated data. D.H., K.C., R.J.C., and J.L.R. wrote the manuscript, and all authors reviewed and approved the final version. D.H. and J.L.R. acquired funding.

## Declaration of Interests

D.H. is an inventor on intellectual property related to chimeric antigen receptor T-cell therapies. All other authors declare no competing interests.

## Notes

https://zenodo.org/records/23397057

