## supplemental-information for "Pan-pediatric brain tumor microRNA networks define immune microenvironment states and responses to CAR T-cell therapy"

### SUPPLEMENTARY INFORMATION

#### Supplementary Tables

**Table S1.** Human tumor and normal brain specimen metadata.

**Table S2.** miRNA differential expression results in tumor vs. tumor-adjacent normal or healthy brain.

**Table S3.** DIPG/DMG and MB differentially expressed miRNA clustering results and immune cell composition correlations.

**Table S4.** Experimentally validated and computationally predicted oncogenic miRNAs (oncoMiRs) and tumor suppressor (TS) miRNAs.

**Table S5.** Gene ontology (GO) enrichment results for DIPG/DMG cluster 1 miRNA and MB cluster 1 miRNA predicted targets.

**Table S6.** RNA differential expression results in tumor vs. tumor-adjacent normal or healthy brain.

**Table S7.** Mouse tumor sample metadata.

**Table S8.** Mouse miRNA and RNA differential expression results in B7H3 CAR T-treated and control CAR-T-treated vs. untreated mice.

**Table S9.** Mouse differentially expressed miRNA clustering results.

**Table S10.** GO enrichment results for upregulated genes in response to CAR T therapy at Day 14 and Day 21 post-treatment.

### Supplementary Figures

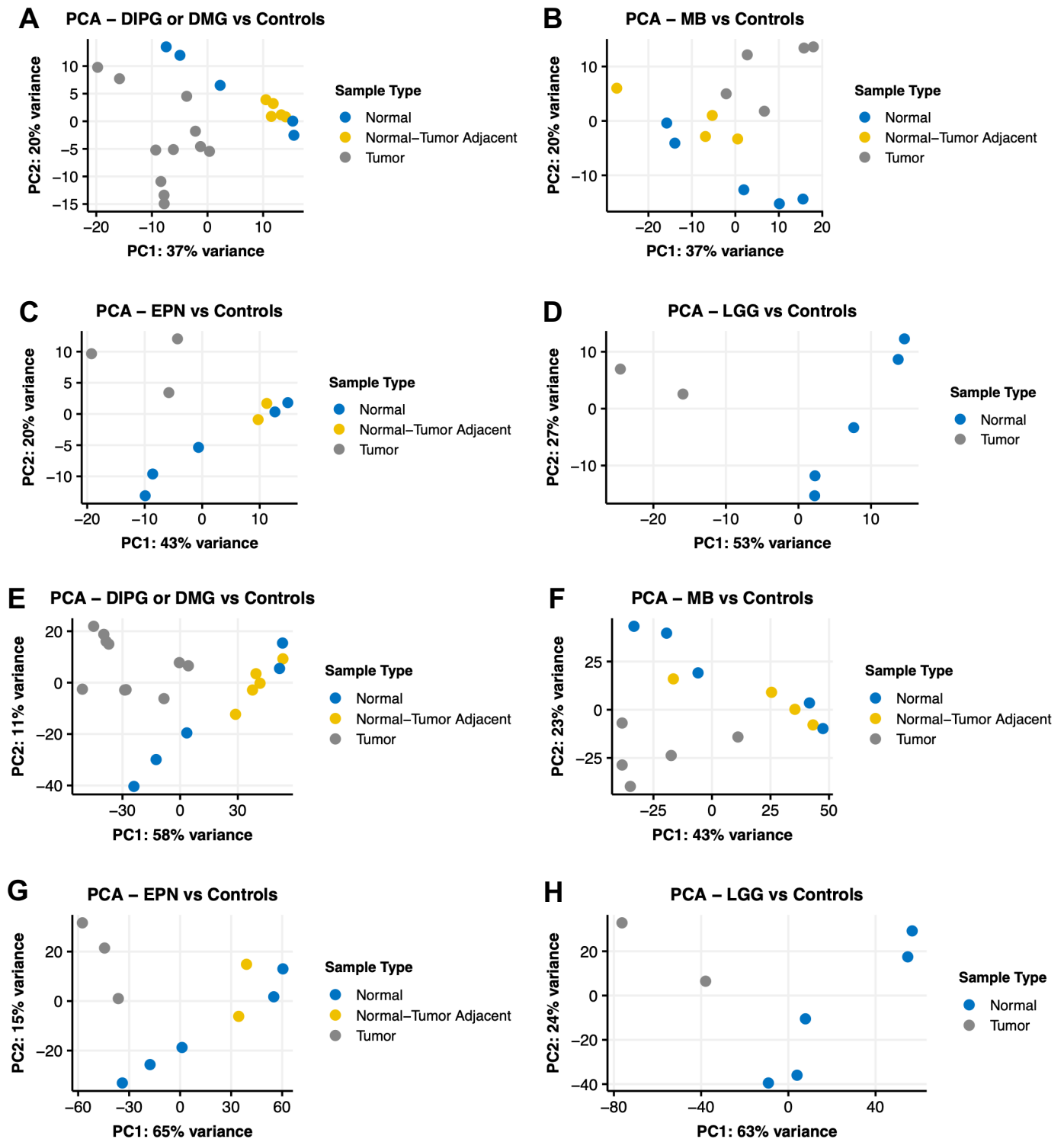

**Figure S1.** Principal component analysis (PCA) plots of CNS tumor and normal brain samples by expression of 500 most variable miRNAs (A-D) and genes (E-H) from small RNA-sequencing and RNA-sequencing, respectively.

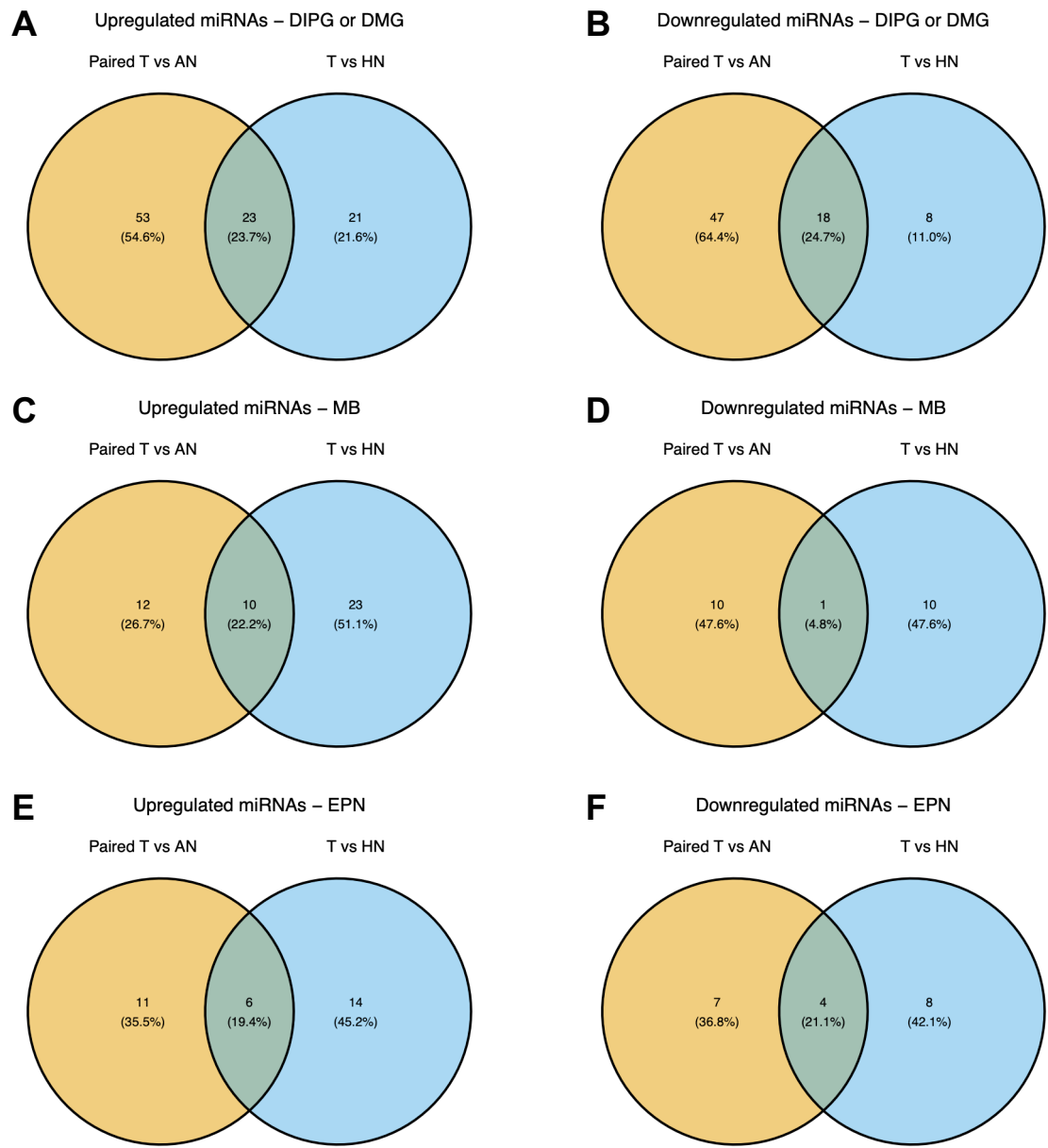

**Figure S2.** **A.** Venn diagram comparing upregulated miRNAs identified in DIPG/DMG versus adjacent normal and versus healthy normal brain. **B.** Venn diagram comparing downregulated miRNAs identified in DIPG/DMG versus adjacent normal and versus healthy normal brain. **C.** Venn diagram comparing upregulated miRNAs identified in DIPG/DMG versus adjacent normal and versus healthy normal brain. **D.** Venn diagram comparing downregulated miRNAs identified in DIPG/DMG versus adjacent normal and versus healthy normal brain. **E.** Venn diagram comparing upregulated miRNAs identified in DIPG/DMG versus adjacent normal and versus healthy normal brain. **F.** Venn diagram comparing downregulated miRNAs identified in DIPG/DMG versus adjacent normal and versus healthy normal brain.

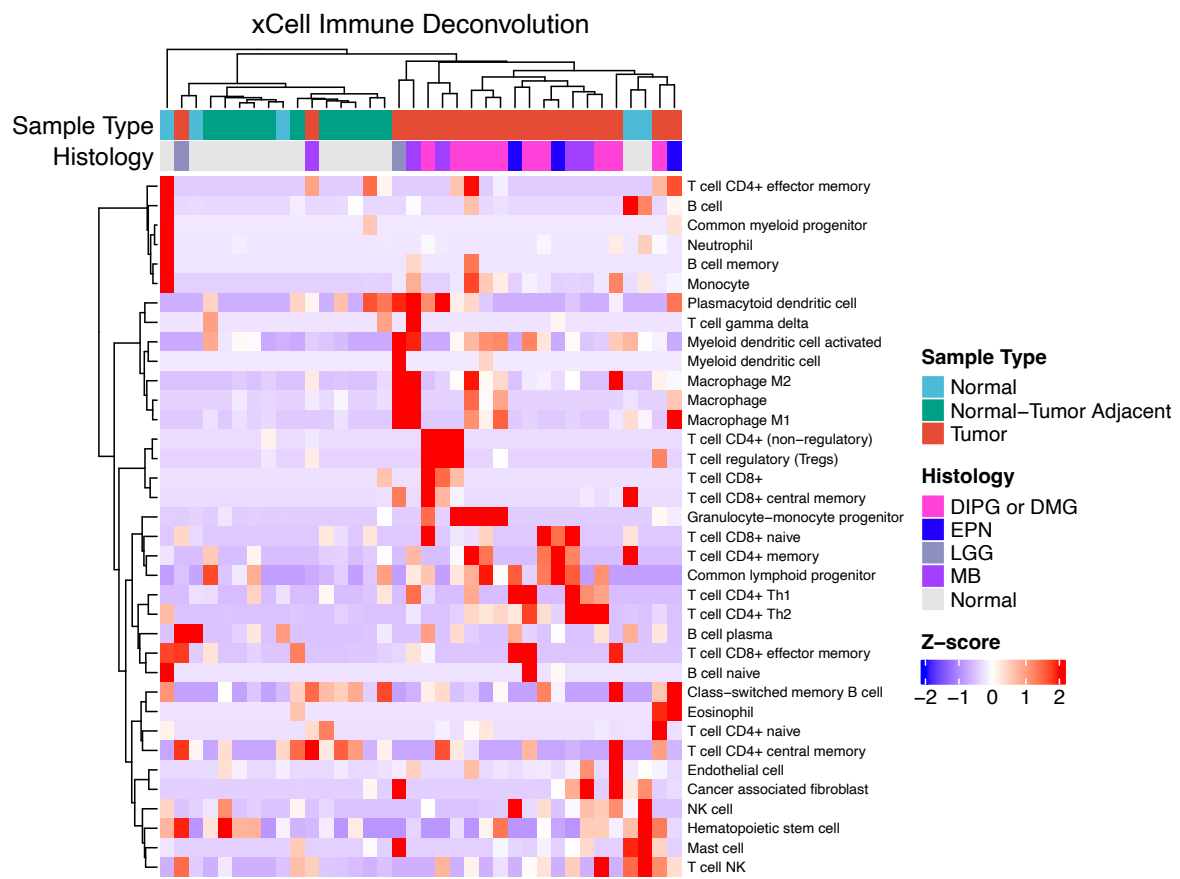

**Figure S3.** Clustered heatmap of XCell-derived immune cell enrichment z-scores in CNS tumor and normal brain samples. Rows and column distances represent Euclidean distances and clustering was performed using the Ward D agglomeration method.

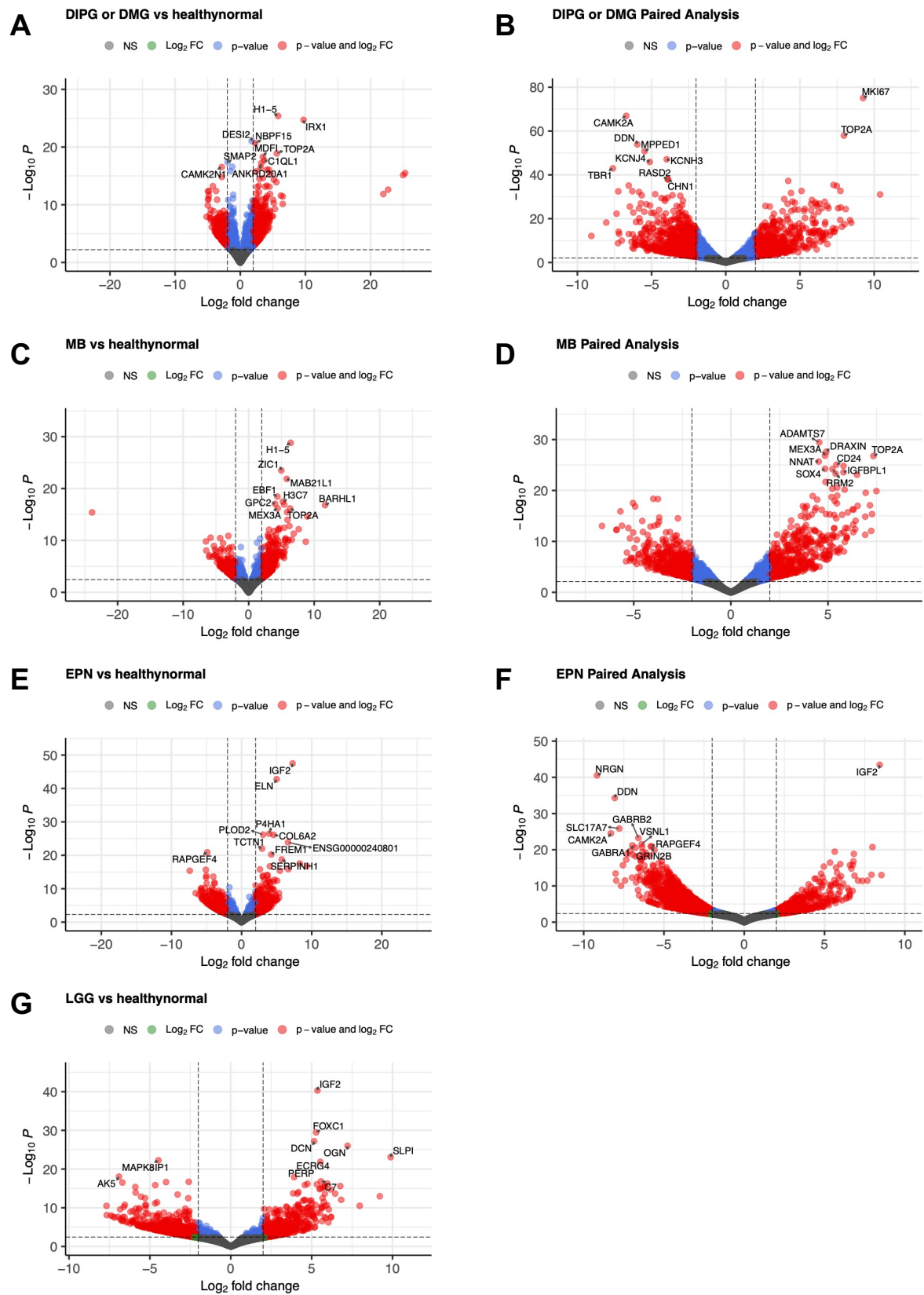

**Figure S4.** Volcano plots displaying log<sub>2</sub>-fold changes and p-values in DIPG/DMG vs. healthy (A) and normal brain (B), MB vs. healthy (C) and normal brain (D), EPN in healthy (E) vs. normal brain (F), and LGG vs. healthy brain (G). Horizontal dashed lines indicate Benjamini-Hochberg adjusted $p < 0.05$, and vertical dashed lines indicate log<sub>2</sub>-fold change $> 2$.

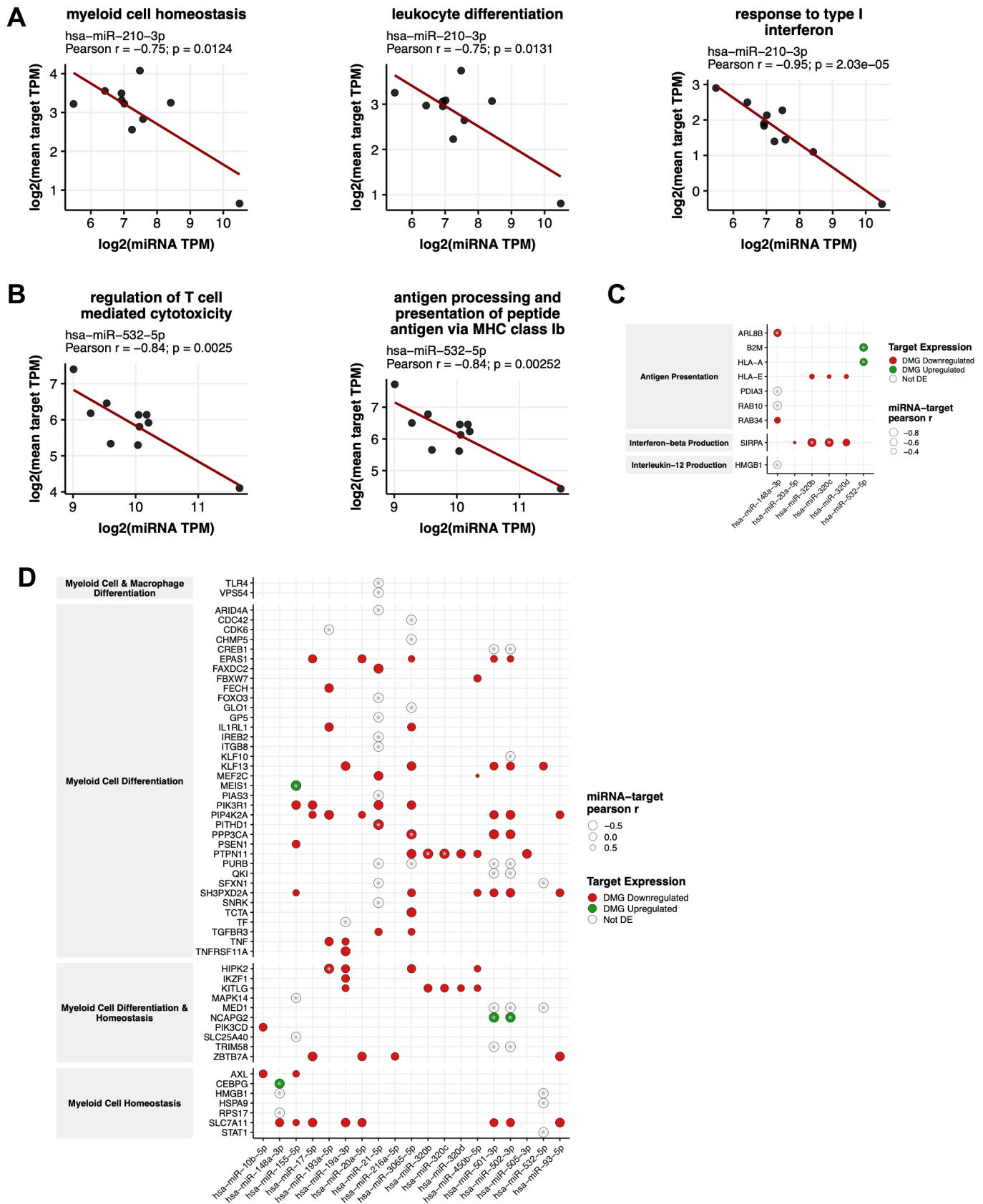

**Figure S5. C.** Dot plot displaying significant interactions (either significant miRNA-target pearson correlation or opposing differential expression) between DIPG/DMG cluster 6 miRNA expression and expression of target genes involved in myeloid cell-related processes

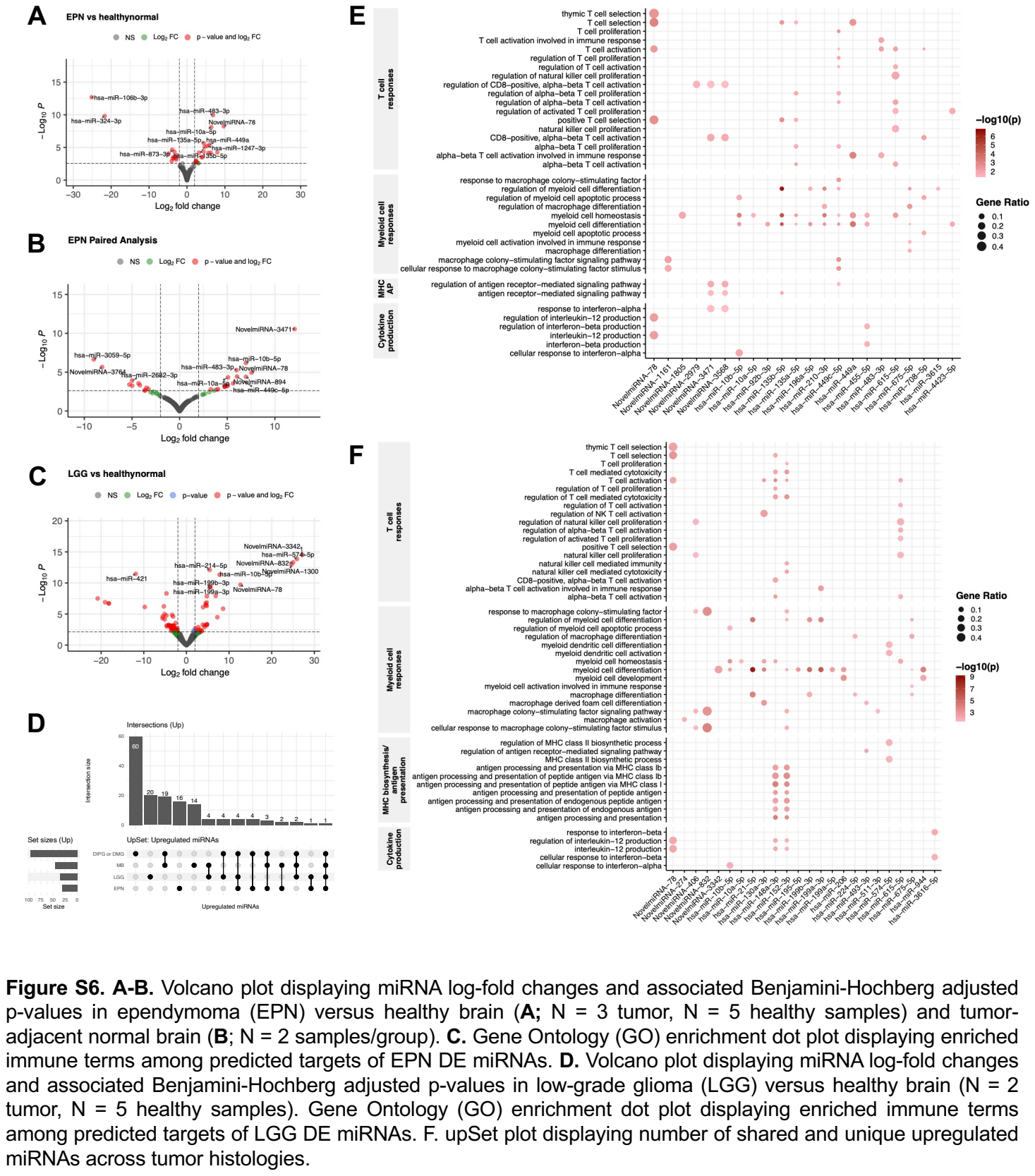

Figure S6. A-B. Volcano plot displaying miRNA log-fold changes and associated Benjamini-Hochberg adjusted p-values in ependymoma (EPN) versus healthy brain (A; N = 3 tumor, N = 5 healthy samples) and tumor-adjacent normal brain (B; N = 2 samples/group). C. Gene Ontology (GO) enrichment dot plot displaying enriched immune terms among predicted targets of EPN DE miRNAs. D. Volcano plot displaying miRNA log-fold changes and associated Benjamini-Hochberg adjusted p-values in low-grade glioma (LGG) versus healthy brain (N = 2 tumor, N = 5 healthy samples). Gene Ontology (GO) enrichment dot plot displaying enriched immune terms among predicted targets of LGG DE miRNAs. F. upSet plot displaying number of shared and unique upregulated miRNAs across tumor histologies.

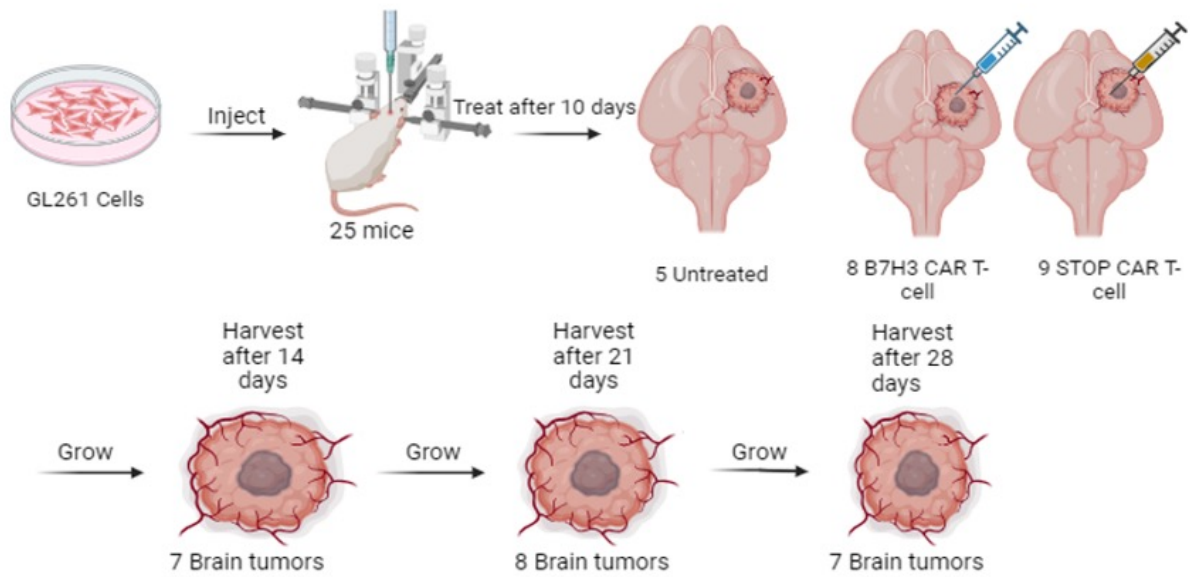

**Figure S7.** Schematic summarizing tumor implantation and subsequent CAR T-cell treatment and tumor harvest method.

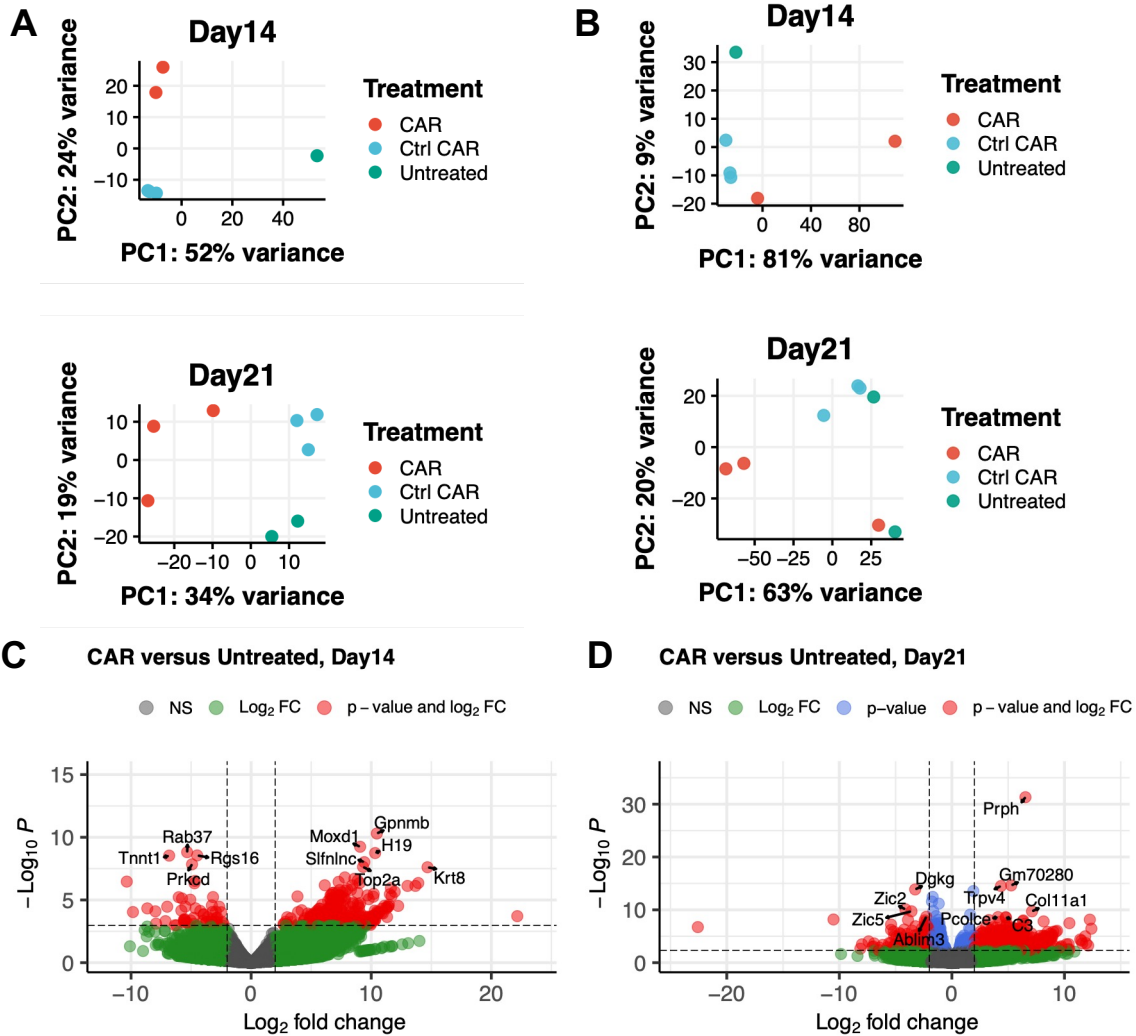

**Figure S8. A-B.** Principal component analysis (PCA) plots of mouse CAR and untreated samples by 500 most variable miRNAs (**A**) and genes (**B**) at Day 14 and Day 21. Volcano plots displaying RNA differential expression in B7H3 CAR T versus untreated mice at Day 14 (**A**) and Day 21 (**B**). N = 2 CAR T-treated and N = 1 untreated at Day 14 and N = 3 CAR T-treated and N = 2 untreated at Day 21.

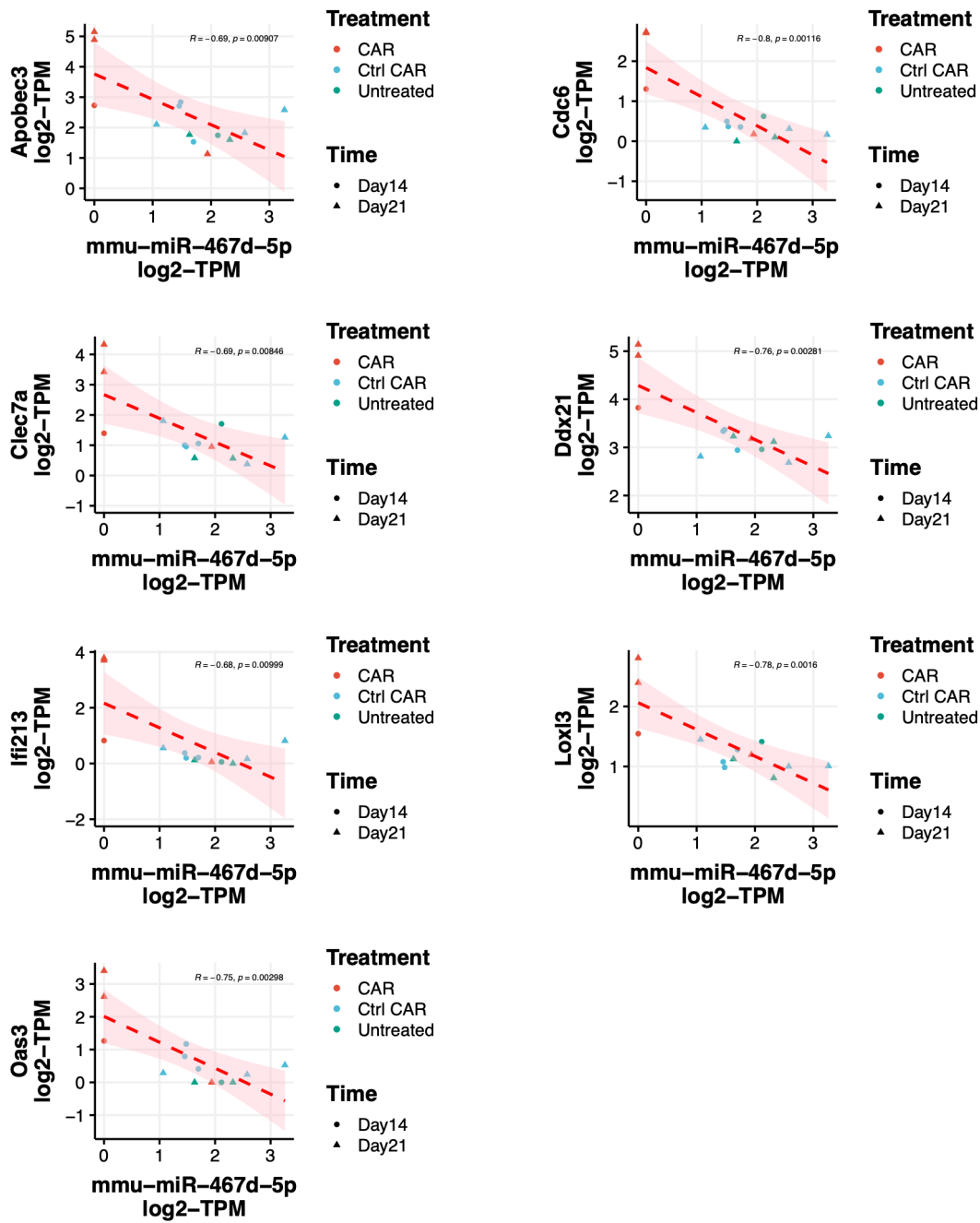

**Figure S9.** Scatter plots of *mmu-miR-467d-5p* expression and expression of target genes *Apobec3*, *Cdc6*, *Clec7a*, *Ddx21*, *Ifi213*, *Loxl3*, *Oas3*, and *Tnfrsf1b*.

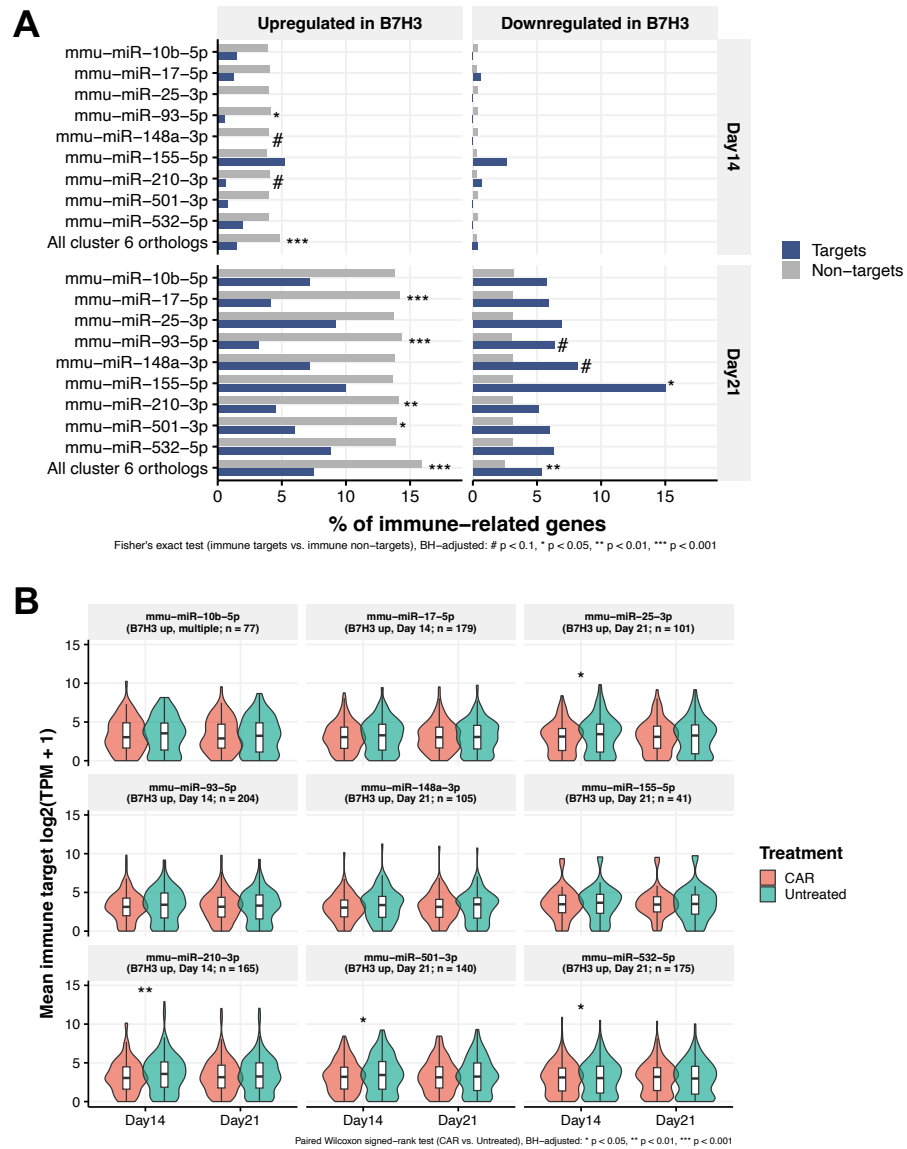

**Figure S10. A.** Bar plots indicating proportion of cluster 6 ortholog targets and non-targets that are up- or downregulated in response to therapy in DMG-bearing mice. Symbols indicate differing thresholds for FDR-adjusted significant odds ratios. **B.** Violin plots of cluster 6 ortholog target expression in response to therapy vs. untreated mice at Day 14 and Day 21. \*\*FDR < 0.05, \*\*\*FDR < 0.01.
